# Recently emerged *Fusarium* chemotypes reprogram wheat defence and detoxification networks during Fusarium head blight development

**DOI:** 10.64898/2026.04.09.717564

**Authors:** Seyedehsanaz Ramezanpour, Nasim Alijanimamaghani, Jason A. McAlister, Ashleigh L. Dale, Stuart J. Cordwell, David Hooker, Jennifer Geddes-McAlister

## Abstract

Fusarium head blight (FHB) is a major threat to global wheat production and food safety due to contamination with mycotoxins, such as deoxynivalenol (DON). The emergence of new mycotoxin chemotypes, including 7-α-hydroxy,15-deacetylcalonectrin (3ANX), presents an evolving challenge for disease management and resistance breeding. Here, we performed a field-based, systems-level proteome analysis of wheat infected with *Fusarium graminearum* strains belonging to the common 15ADON and recently emerged 15ADON/3ANX chemotypes. Across host and pathogen, we quantified more than 9,200 proteins, providing extensive coverage of infection-associated molecular responses. Infection with 15ADON/3ANX strains suppressed canonical wheat detoxification pathways while promoting structural and oxidative defence responses. Concurrently, the fungal proteome of 15ADON/3ANX-producing strains indicated altered mitochondrial ribosome function and alternative virulence strategies. Further investigation of the host-pathogen interface defined hub protein networks negatively regulating classical detoxification markers, suggesting coordinated regulation of host defence responses regardless of chemotype. Molecular responses were linked to field phenotypes by quantification of DON-3-glucoside/DON ratios and disease severity, defining positive correlations in 15ADON infections, which were abolished upon 15ADON/3ANX infection, indicating chemotype-specific evasion or suppression of host defenses. These findings demonstrate reprogramming of host–pathogen interaction networks and reveal molecular targets that may inform chemotype-aware breeding strategies to enhance crop resilience.

## Introduction

Fusarium head blight (FHB), caused predominantly by *Fusarium graminearum*, is a globally widespread disease of wheat (*Triticum aestivum*) that severely threatens both plant yield and grain quality (Fernando *et al*., 2021). The economic and agronomic effects of FHB threaten food safety as infected grains accumulate trichothecene mycotoxins, such as deoxynivalenol (DON) and its acetylated derivatives: 15-acetyldeoxynivalenol (15ADON) and 3-acetyldeoxynivalenol (3ADON) (Audenaert *et al*., 2013; Varga *et al*., 2015). These mycotoxins inhibit eukaryotic protein synthesis, disrupt cellular metabolism, and contribute directly to fungal virulence (Nishiuchi *et al*., 2006; Perochon & Doohan, 2024), making them crucial factors in the pathogenesis of FHB and a significant driver for disease resistance breeding strategies. Critically, the emergence of new chemotypes producing 7-α hydroxy,15-deacetylcalonectrin (3ANX) is becoming more widespread and prominent in North America and Europe, highlighting challenges associated with evasion of disease resistance and effective detoxification mechanisms (Foroud *et al*., 2019; Pierron *et al*., 2022; Schiwek *et al*., 2022; Laraba *et al*., 2023). The shifting chemotype profile underscores the need to decode wheat’s molecular defence and detoxification responses in a chemotype-specific manner to better understand how to combat these emerging threats and maintain mycotoxin detection and tolerance levels and ensure food safety.

FHB resistance in wheat is complex and polygenic, involving divergent molecular pathways with resistance categorized into: Type I (resistance to initial infection), Type II (resistance to spread within the spike), Type III (resistance to mycotoxin accumulation), Type IV (resistance to kernel damage), and Type V (tolerance to yield loss) (Ma *et al*., 2025). Recent multi-omics approaches have begun to unravel the molecular foundations of these resistance types, integrating transcriptomic, metabolomic, and proteomic data to identify key regulatory pathways and candidate resistance genes (Dhokane *et al*., 2016; Wu *et al*., 2022; Sirangelo, 2024). Notably, proteomics has developed as a powerful tool for dissecting plant–pathogen interactions, through high-resolution understanding of protein abundance, post-translational modifications, and protein–protein interactions (Ding *et al*., 2017; Liu *et al*., 2022b; Kosová *et al*., 2023). In wheat, proteomics has identified several defence- and detoxification-related proteins with differential production during FHB infection, including pathogenesis-related (PR) proteins, glutathione S-transferases (GSTs), glycosyltransferase, ATP-binding cassette (ABC) transporters, and chitinases (Walter *et al*., 2015; Yang *et al*., 2021; Buchanan *et al*., 2025). These proteins play significant roles in detoxification, signal transduction, and structural defence, and their production patterns often vary among chemotypes (Kazan & Gardiner, 2018).

The chemotype-specific basis of wheat defence and detoxification suggests a customized molecular approach that adapts to the virulence profile of the pathogen. For instance, 3ADON-producing strains may exhibit elevated but variable aggressiveness of DON accumulation compared to 15ADON strains under specific field conditions, with complementary transcriptomic studies demonstrating enhanced regulation of virulence factors and tailored host responses (Puri *et al*., 2016). Specifically, recently emerged NX and 15ADON/3ANX chemotypes that produce novel trichothecenes may activate new defence pathways or suppress conventional resistance mechanisms. For instance, we demonstrated that wheat inoculated with 15ADON/3ANX-producing strains exhibited limited production of GSTs as classical defence mechanisms, which was accompanied by the exclusive production of structural and oxidoreductase-related proteins, showing a shift toward alternate defence mechanisms (Ramezanpour *et al*., 2026). Understanding these interactions is critical for discovering new biomarkers of resistance and developing chemotype-specific breeding methods for improved resilience. Conversely, beyond the host, fungal adaptation strategies also contribute to chemotype-specific pathogenicity. For example, modulation of mitochondrial function, ribosomal biogenesis, and secondary metabolite biosynthesis enables *F. graminearum* to optimize its virulence profile under varying environmental and host conditions (Atanasova-Penichon *et al*., 2016; Song *et al*., 2024). Protein mapping have revealed that these adaptations are reflected in the production of ergosterol biosynthesis enzymes, proteasome-associated proteins, and oxidative stress regulators, particularly in 3ANX-producing strains (Song *et al*., 2024; Ramezanpour *et al*., 2026). These findings imply that chemotype evolution is not only a modulation of plant response pathways but also a broader reprogramming of fungal metabolism and pathogenicity.

In this study, we leveraged high-resolution mass spectrometry (MS) using data-independent acquisition (DIA) to conduct a comprehensive proteomics investigation of wheat infected under field conditions with *F. graminearum* strains that produce either 15ADON or a combination of 15ADON and 3ANX. We identified over 9,200 proteins originating from both host and pathogen and representing a 1.8-fold increase in protein identifications compared to our previous data-dependent acquisition (DDA) assessments. Our in-depth profiling demonstrate that the presence of 3ANX fundamentally reshapes classical DON-associated defense pathways; specifically, 15ADON/3ANX infections resulted in reduced abundance of canonical detoxifying proteins, such as GSTs, and a shift toward alternative structural and oxidative defense mechanisms, including chitinases, defensins, heat shock proteins, and glycosyltransferases. From the fungal perspective, 498 proteins were found to be exclusive to the 15ADON/3ANX chemotype, involving mitochondrial ribosome function and virulence-associated mechanisms (e.g., cell wall integrity, oxidative stress response, plant immunity suppression), supporting alternative chemotype-specific mechanisms to counteract or evade plant defenses. Through network analyses, we identified three wheat hub proteins (i.e., DSK2a, elongation factor 1-γ2, and uncharacterized) that exhibited inverse correlation profiles to standard detoxification proteins, suggesting their role in coordinating these altered responses regardless of chemotype. To correlate molecular observations with field phenotypes, we measured the ratio of D3G (less toxic) to DON, as well as disease severity and mapped the values across all samples. Here, we observed that the positive correlation between detoxification activity and disease severity observed in 15ADON infections was lost in the presence of 15ADON/3ANX, indicating a disruption of traditional resistance-linked outcomes. Collectively, these findings provide a high-resolution molecular framework for understanding how recently emerged chemotypes bypass established wheat defenses and highlight novel targets for chemotype-specific precision breeding and agriculture through resistance breeding for improved global food security and safety.

## Materials and Methods

### Fungal culture and macroconidia preparation

*Fusarium graminearum* strains (Table S1) were prepared as previously described (Tamburic-Ilincic *et al*., 2007; Ramezanpour *et al*., 2026) with some modifications. Briefly, ten 1 cm² agar plugs from each actively growing culture were transferred into Bilay’s liquid medium (per liter: 8.0 g KNO□, 4.0 g KH□PO□, 2.0 g KCl, 2.0 g MgCl□, 2.0 g sucrose plus 2 mL of a solution containing 20 mg per 100 mL of FeCl_3_, MnSO_4_, and ZnSO_4_). For each strain, three flasks were inoculated and one flask served as an uninoculated control. Cultures were grown for 21 days at 25□°C on a rotary shaker at 120 rpm. Strains were diluted to 5 × 10□ spores/mL with four technical replicates were generated by dividing the macroconidial suspension or mycelial filtrate into four sterile containers, each used as an independent inoculum source.

### Wheat cultivation and inoculation

The misted field experiment included 20 *F. graminearum* strains, and one uninoculated control, each inoculated in four replicates arranged in a split-plot design (main-plot for disease severity, sub-plot for proteomic samples) as previously described (Ramezanpour *et al*., 2026). The winter wheat cultivar 25R40 (susceptible to FHB (Xia *et al*., 2021)), distributed by Pioneer (Guelph, ON, Canada), was planted in October 2022 at the Ridgetown Research Station, University of Guelph (Guelph, ON, Canada). Average temperatures and rainfall during the post-inoculation period (June-July 2023) was 23.8 °C and 3.88 mm, respectively, which is conducive to FHB development. Point inoculations were performed at 50% anthesis (Dyer *et al*., 2005; Geddes *et al*., 2008) in June 2023 using 100 µL of standardized macroconidia suspensions of *F. graminearum* or a media control. Following inoculation, plots were subjected to overhead mist irrigation for 21 days post-inoculation (dpi) to promote FHB development and harvested with 10 heads per strain pooled into a single collection bag. Samples were transported on ice and stored at −80 °C until protein extraction.

### Protein extraction

Wheat heads were treated as previously described (Liu *et al*., 2022a; Ramezanpour *et al*., 2026). In brief, wheat spikes were ground in liquid nitrogen with a mortar and pestle and resuspended in a lysis buffer (100 mM Tris-HCl, pH 8.5) followed by reduction (dithiothreitol; 10 mM), alkylation (iodoacetamide; 5 mM), and acetone precipitation (-20 °C). Precipitated proteins were resuspended in 8 M urea/40 mM HEPES, quantified (Wiśniewski *et al*., 2009), and digested with trypsin/LysC (50:2, protein: enzyme with 0.5 µg/µL enzyme mix). Peptides were purified using STop And Go Extraction Tips (Rappsilber *et al*., 2007).

### Liquid chromatography – tandem mass spectrometry (LC-MS/MS) of peptides

Analysis of peptides was performed on a Thermo Scientific™ Orbitrap Exploris 480™ instrument coupled online to a Thermo Scientific™ Vanquish Neo™ UHPLC system equipped with a 75 μm × 35 cm home-packed C18 resin (1.9 μm) column. Purified peptides were re-constituted in 5% acetonitrile (MeCN), 0.1% formic acid (FA) and separated with reverse phase (RP) chromatography by increasing the concentration of the mobile phase over a 75-min gradient from 97% buffer A (0.1% [v/v] FA), 3% buffer B (80% [v/v] MeCN, 0.1% [v/v] FA) to 98% buffer B, 2% buffer A at a flow rate of 300 nL/min. The mass spectrometer was operated in DIA mode. The instrument was configured to perform a survey MS1 scan at 30,000 resolution, scan range 350-1650 mass-to-charge (*m/z*), and maximum injection time of 54 ms. Precursors were fragmented using stepped higher-energy collisional dissociation (25%, 27%, 30%) and analysed using 20 variable isolation windows over the scan range 350-1650 *m/z*. MS/MS scans were performed at a resolution of 30,000 and a normalised automatic gain control (AGC) target of 1000%.

### LC-MS/MS data processing

Raw MS data were processed using Spectronaut (version 20.3.251119.92449). Spectra were searched against *Triticum aestivum* (Sept. 2025, 17,234 sequences) and *Gibberella zeae* (Sept. 2025, 14,146 sequences) UniProt databases. Search parameters included: trypsin specificity with up to two missed cleavages; carbamidomethylation of cysteine as a fixed modification; methionine oxidation and N-terminal acetylation as variable modifications. Peptide and protein identifications were filtered at a 1% false discovery rate (FDR). Mass tolerance was set to 4.5 ppm for precursor ions and 20 ppm for fragment ions.

### Bioinformatics and statistical analysis

Data processing and visualization were performed with Perseus (version 1.6.2.2) and ProteoPlotter (Tyanova *et al*., 2016; Olabisi-Adeniyi *et al*., 2025). Data were filtered for protein identifications by at least two peptides and intensities were log2-transformed. Valid value filtering was conducted using proteins found in at least three of four replicates in at least one group (standard analysis) or within each group (core proteome analysis), followed by imputation with a downshift of 1.8 and a breadth of 0.3 standard deviations. Data were visualized using principal component analysis (PCA), dynamic range plots with protein intensities constructed for each sample, and 1D annotation enrichment in protein abundance across defined protein annotations using Student’s t-test, p-value < 0.05; FDR = 0.05, and score <-0.5, >0.5 (Benjamini & Hochberg, 1995; Cox & Mann, 2012). Proteins were annotated according to Gene Ontology (GO), Biological Processes (GOBP), GO Molecular Function (GOMF), GO Cellular Component (GOCC), and UniProt Keywords. GraphPad Prism (v10) was used to construct box plots, and statistical testing was performed using ANOVA, Dunnett test with a p-value < 0.05. For fungal proteins, strains were grouped by chemotype (15ADON vs. 15ADON/3ANX), and protein abundances were compared between groups using unpaired t-tests (GraphPad Prism v10) with a p-value < 0.05 considered significant. Protein-protein interaction networks for wheat and fungal core proteins were generated using the STRING database (high confidence = 0.7) (Szklarczyk *et al*., 2019); the networks were reconstructed in Cytoscape. The Cyto-Hubba plugin in Cytoscape software was used to identify hub proteins (20 nodes with the highest interaction) in the network using 12 provided CytoHubba calculation algorithms, including Degree, Edge Percolated Component (EPC), Maximum Neighborhood Component (MNC), Density of Maximum Neighborhood Component (DMNC), Maximal Clique Centrality (MCC), Bottleneck, Closeness, Betweenness, Stress, Radiality, Eccentricity, and Clustering Coefficient.

### Mycotoxin quantification and disease severity

At 21 dpi, wheat heads were air dried, threshed, homogenized, and weighed as previously described (Limay-Rios & Schaafsma, 2021). Mycotoxin standards were obtained from Biopure™ standards (Romer Labs®, Tulln, Austria). Mycotoxin extraction was performed as previously described by high performance liquid chromatography coupled with MS on a QTRAP 6500+ hybrid triple quadrupole/linear ion trap system over a 22 min gradient (Limay-Rios & Schaafsma, 2021). DON and D3G concentrations were measured for each strain and used to calculate the D3G/DON ratio as an indicator of DON glycosylation efficiency. Disease severity scores were incorporated as a quantitative measure of chemotype specific disease impact. The percentage of disease severity was calculated based on the number of spikelets with disease symptoms (e.g., bleaching, water-soaked spots, shriveled) divided by the total number of spikelets in each head (the average of ten infected heads in each plot). Pearson correlation analyses were performed between detoxification associated proteins, the D3G/DON ratio, and severity for wheat inoculated with 15ADON and 15ADON/3ANX producing strains. Correlation matrices were generated in GraphPad Prism (v10) to compare chemotype dependent trends across detoxification pathways, DON conversion, and disease severity.

## Results

### Proteomic profiling via DIA-MS provides enhanced depth of proteome coverage

To investigate variations in wheat response to *F. graminearum* strains producing 15ADON or the combination of 15ADON/3ANX, we reproduced FHB infection in the field using point inoculation with fungal mycelium or spores (Fig. 1A; Table S1). Following three-weeks of field growth, samples were collected and high-resolution MS-based proteomics was performed on the samples using DIA to systematically fragmenting all peptides in a given *m/z* range allowing unbiased acquisition rather than selecting a pre-specified number of abundant peptides (as per DDA) (Fröhlich *et al*., 2024). We identified 9,248 proteins after valid value filtering (proteins identified in 3 of 4 replicates) with 5,481 designated as wheat proteins and 3,766 designated as fungal proteins (Fig. 1B). Notably, DIA-MS provided increased coverage of the wheat and fungal proteome by 1.4- and 2.7-fold, respectively, compared to our previous study using DDA-MS (Ramezanpour *et al*., 2026), supporting a greater depth of proteome detection for both biological systems with a notable improved depth of information for the pathogen.

**Figure 1.**
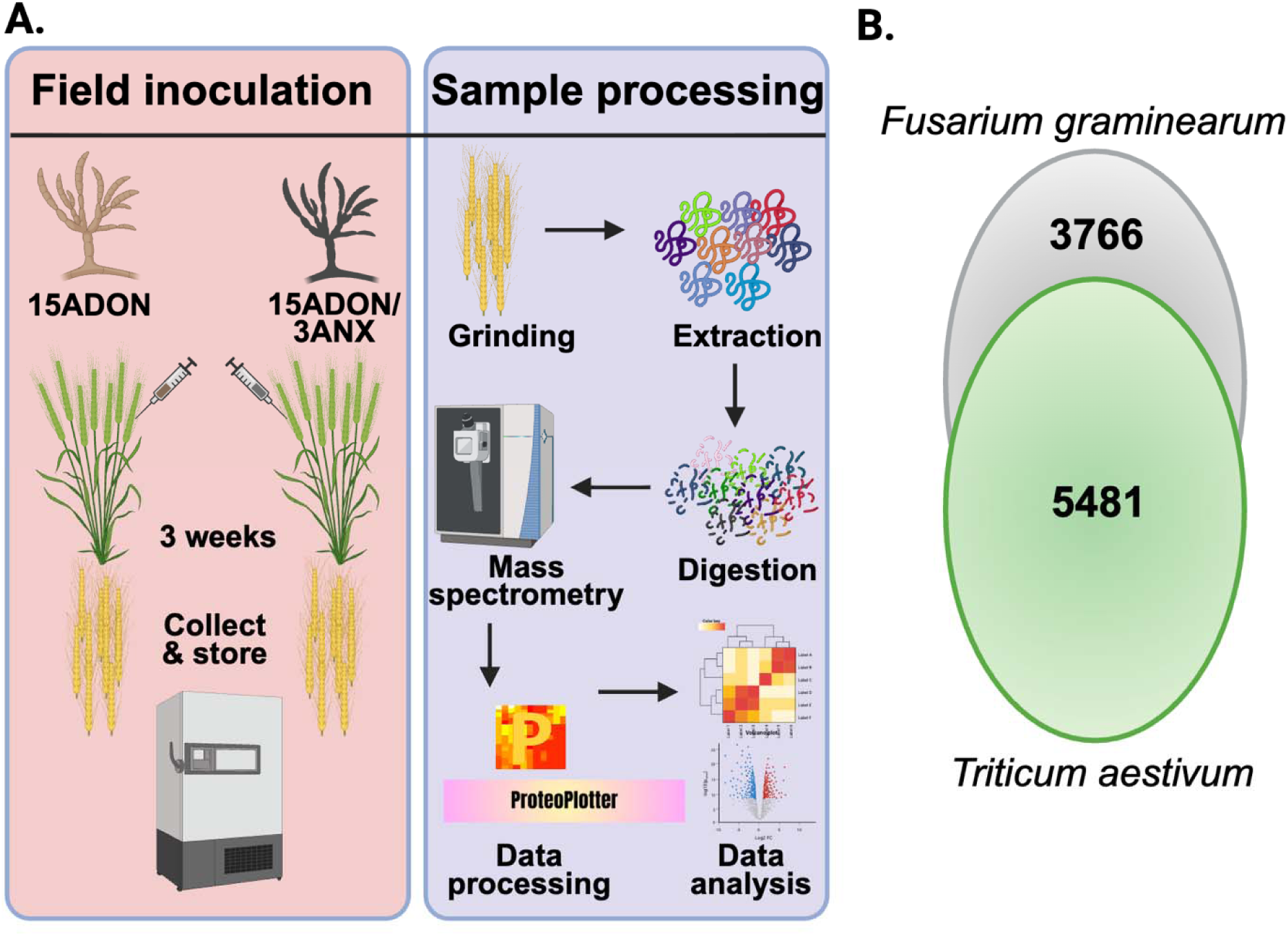
Overview of field inoculation strategy and DIA-MS-based proteomics workflow. **A.** Wheat plants were point-inoculated in the field with *Fusarium graminearum* strains representing the 15ADON and 15ADON/3ANX chemotypes, alongside an uninoculated control. Following a three-week infection period, wheat heads were collected, and proteins were extracted, digested, and purified for data-independent acquisition (DIA) mass spectrometry (MS). **B.** Total number of identified proteins (detected in ≥3 of 4 biological replicates) assigned to *Triticum aestivum* (green) and *F. graminearum* (grey). Each biological replicate consisted of a pooled sample of ten wheat heads, and inoculum for each fungal strain was prepared from four independent biological cultures.

**Table 1.**
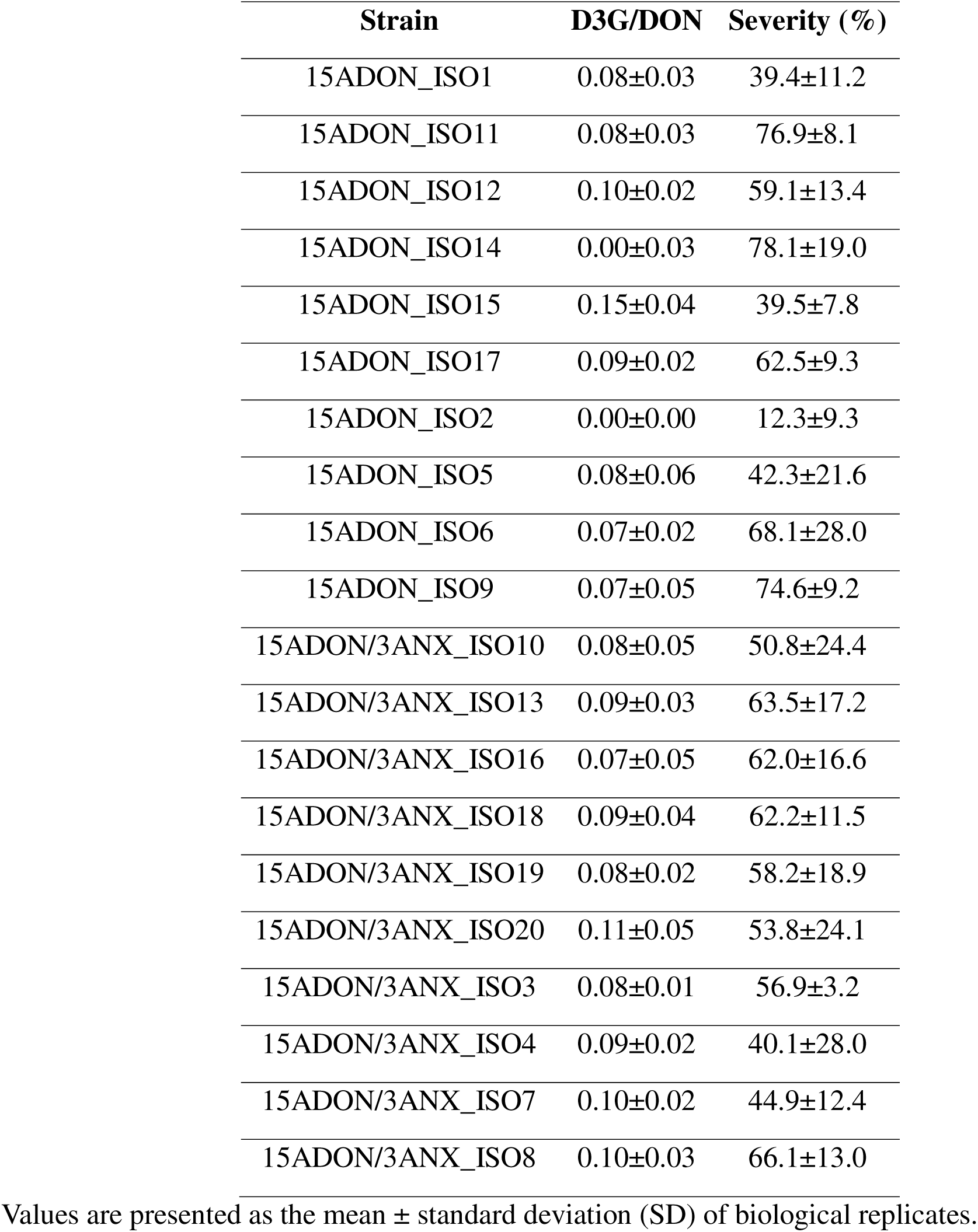
D3G/DON ratio and disease severity (%) in wheat inoculated with 15ADON and 15ADON/3ANX-producing *F. graminearum* strains.

| Strain | D3G/DON | Severity (%) |
| --- | --- | --- |
| 15ADON_ISO1 | 0.08±0.03 | 39.4±11.2 |
| 15ADON_ISO11 | 0.08±0.03 | 76.9±8.1 |
| 15ADON_ISO12 | 0.10±0.02 | 59.1±13.4 |
| 15ADON_ISO14 | 0.00±0.03 | 78.1±19.0 |
| 15ADON_ISO15 | 0.15±0.04 | 39.5±7.8 |
| 15ADON_ISO17 | 0.09±0.02 | 62.5±9.3 |
| 15ADON_ISO2 | 0.00±0.00 | 12.3±9.3 |
| 15ADON_ISO5 | 0.08±0.06 | 42.3±21.6 |
| 15ADON_ISO6 | 0.07±0.02 | 68.1±28.0 |
| 15ADON_ISO9 | 0.07±0.05 | 74.6±9.2 |
| 15ADON/3ANX_ISO10 | 0.08±0.05 | 50.8±24.4 |
| 15ADON/3ANX_ISO13 | 0.09±0.03 | 63.5±17.2 |
| 15ADON/3ANX_ISO16 | 0.07±0.05 | 62.0±16.6 |
| 15ADON/3ANX_ISO18 | 0.09±0.04 | 62.2±11.5 |
| 15ADON/3ANX_ISO19 | 0.08±0.02 | 58.2±18.9 |
| 15ADON/3ANX_ISO20 | 0.11±0.05 | 53.8±24.1 |
| 15ADON/3ANX_ISO3 | 0.08±0.01 | 56.9±3.2 |
| 15ADON/3ANX_ISO4 | 0.09±0.02 | 40.1±28.0 |
| 15ADON/3ANX_ISO7 | 0.10±0.02 | 44.9±12.4 |
| 15ADON/3ANX_ISO8 | 0.10±0.03 | 66.1±13.0 |

Values are presented as the mean ± standard deviation (SD) of biological replicates.

### Distinct proteome signatures in wheat reveal altered defense strategies against 15ADON and emerging 3ANX mycotoxins

Based on *F. graminearum* strains producing 3ANX with enhanced prevalence and virulence relative to DON (Varga *et al*., 2015; Pierron *et al*., 2022), we sought to examine changes in wheat response based on mycotoxin chemotype. For the initial analysis, we focused on the core wheat proteome, which consisted of 2,773 proteins identified in at least three of four biological replicates within each sample group. A principal component analysis (PCA) revealed moderate separation by samples along component 1 (28%) and treatment for component 2 (18.1%); however, clear clustering within treatments or conditions was not evident (Fig. 2A). Next, to examine replicate reproducibility, mean Pearson correlation values were calculated at 86.8% for control samples, 88.5% for 15ADON-treated samples, and 86.3% for 15ADON/3ANX-treated samples, indicating strong correlation among replicates in each treatment group from samples gathered from field trials (Fig. S1). The dynamic range of protein intensities was assessed across the conditions, spanning four orders of magnitude for control, 15ADON, and 15ADON/3ANX samples, suggesting strong coverage of the proteome from high to low abundance proteins (Fig. 2B).

**Figure 2.**
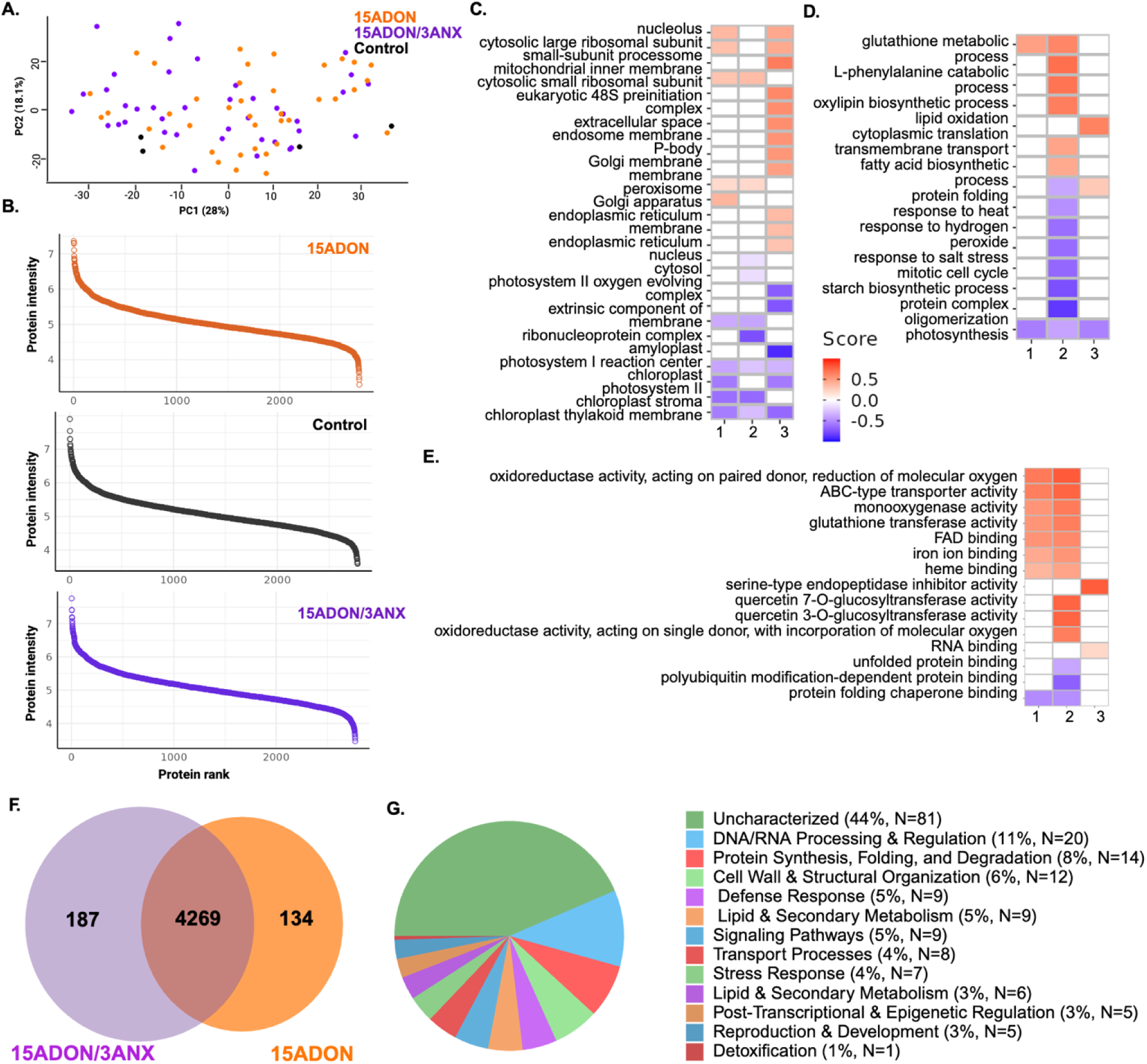
Chemotype□dependent restructuring of the core wheat proteome. **A.** Principal component analysis. **B.** Dynamic range plots (log10 y-axis) showing the distribution of protein intensities across ranked proteins for control (black), 15ADON (orange), and 15ADON/3ANX (purple) conditions. **C–E.** 1D annotation enrichment analyses highlighting chemotype driven shifts in protein localization (GO Cellular Compartment), biological activity (GO Biological Processes), and molecular functions (GO Molecular Function). Enrichment was assessed using Student’s t test (p < 0.05) with FDR correction (0.05) and significance thresholds of score < –0.5 or > 0.5. Comparisons: (1) 15ADON/3ANX vs. control, (2) 15ADON vs. control, (3) 15ADON/3ANX vs. 15ADON; Red = enriched in first factor of comparison, Blue = enrichment in second factor of comparison. **F.** Venn diagram showing the overlap of wheat proteins detected across chemotype treatments, including proteins shared between both treatments (4,269) and those uniquely identified in 15ADON (134) or 15ADON/3ANX (187) samples. **G.** Functional categorization (GOBP) of proteins exclusive to the 15ADON/3ANX treatment, illustrating enrichment of structural, metabolic, and defense associated processes.

Next, to evaluate enrichment of protein annotations across Gene Ontology categories, we performed a 1D annotation enrichment of wheat proteins upon comparison of chemotypes vs. control and between chemotypes (Cox & Mann, 2012). Here, by GOCC, we observed enrichment of membrane and mitochondrial inner membrane as a function of chemotype treatment, compared to enrichment within the control of proteins associated with the chloroplast, chloroplast stroma, chloroplast thylakoid membrane, and the ribonucleoprotein complex, suggesting enhanced energy production and reduced photosynthesis-associated functions upon infection (Fig. 2C). We also observed substantial remodeling upon comparison of chemotypes in the presence of 3ANX impacting many translation-associated functions (e.g., endoplasmic reticulum, nucleus). Additionally, analysis by GOBP showed enrichment of glutathione metabolic process across both chemotypes relative to the control, and a reduction of photosynthesis in infected vs control, as well as in 15ADON/3ANX vs 15ADON (Fig. 2D). Notably, the 15ADON treatment relative to the control led to the enrichment of proteins associated with diverse functions, including the phenylalanine catabolic pathway, oxylipin biosynthetic process, lipid oxidation, transmembrane transport, and fatty acid biosynthetic process, along with a reduction of abundance for proteins associated with stress response, protein folding, and starch biosynthesis, indicating that the 15ADON/3ANX chemotype masks responses of these categories within wheat. Chemotype comparisons revealed enrichment of cytoplasmic translation and protein folding in the presence of 3ANX and reduced photosynthesis. Finally, based on GOMF, we observed consistent enrichment between chemotypes for proteins associated with oxidative stress, phenylpropanoid biosynthesis, detoxification (ABC transporters, monooxygenases, glutathione transferases), and ion binding (e.g., FAD, iron, heme), and a reduction in protein folding compared to control (Fig. 2E). Exposure to 15ADON showed unique enrichment for quercetin 7- or 3-O-glucosyltransferase activity, suggesting that presence of 3ANX does not initiate this response, as well as additional oxidoreductase enrichment. Between chemotypes, we observed enrichment of serine-type endopeptidase inhibitor binding and RNA binding in the presence of 3ANX. Together, these assessments of enrichment across GO categories demonstrates activation of anticipated functions, such as oxidoreductase and glutathione activities, and reduced photosynthesis in the presence of the 15ADON and 15ADON/3ANX chemotypes, as well as distinct patterns of suppressed enrichment of host defense response in the presence of 3ANX.

### Chemotype-dependent host protein abundance highlights 3ANX-specific defense mechanisms

Given our observation of altered wheat protein production profiles dependent upon mycotoxin chemotype treatment across all host samples, we aimed to identify wheat proteins exclusively detected in the presence of 3ANX. Here, we observed 4,269 host proteins shared between 15ADON/3ANX and 15ADON treatment, with 187 proteins exclusive to 15ADON/3ANX and 134 proteins exclusive to 15ADON treatment (Fig. 2F). We focused on the 3ANX-exclusive proteins and observed that almost 50% were uncharacterized proteins with diverse functions (Fig. 2G; Table S2). Proteins associated with functions in defence and stress response accounted for 9% of the proteins identified, including chitinase, defensins, and heat shock proteins (e.g., HSP26), which may limit *Fusarium* colonization and mycotoxin damage (Cheng *et al*., 2015; Leannec-Rialland *et al*., 2021; Shafikova *et al*., 2024). Proteins associated with transport, signalling pathways, and detoxification were also identified, including glycosyltransferases, with defined roles in mycotoxin detoxification (He *et al*., 2018, 2020). These findings identify wheat’s response in the presence of 3ANX, including established responses to mycotoxin contamination and defence response, as well as potential novel protein activation mechanisms.

### Core wheat proteome analysis identifies conserved defense in wheat-pathogen interactions across diverse F. graminearum strains

Given the differences in wheat proteome remodeling observed across mycotoxin chemotypes, and the differential strain sources (e.g., mycelium or spore, wheat or corn, collections; Table S1), we investigated strain-specific responses from the wheat perspective. Among the core proteome, we prioritized 32 proteins with defined roles in pathogenesis-related response (e.g., PR proteins), defence response (e.g., chitinases, glucanases), and mycotoxin defence (e.g., glutathione transferases) (Jain & Khurana, 2018) (Table S3). For PR proteins and proteins associated with reactive oxygen species protection (e.g., catalase, peroxidase, superoxide dismutase), we did not observe any significant changes in abundance across the strains compared to the untreated control, suggesting that these general plant defenses are no longer dominant by 21 dpi (Fig. S2). We observed similar trends across the identified catalase (N=4), peroxidase (N=19) and superoxide dismutase (N=1) with fluctuations in enzyme abundance but not at a significant level.

Conversely, an assessment of general defense responses, including activation of lignin biosynthesis process (cinnamyl alcohol dehydrogenase) (Fig. 3A), defence response to fungus (thaumatin-like protein) (Fig. 3B), and signal transduction (14-3-3 domain-containing protein and leucine-rich repeat-containing *N*-terminal plant-type domain-containing protein) (Fig. 3C) showed significant changes in abundance across infection with different fungal strains compared to the untreated control. Notably, a single 15ADON-producing strain (#6) resulted in significantly elevated abundance of cinnamyl alcohol dehydrogenases. Despite limited differences in defense-associated proteins across the strains, we identified significant increases in abundance of proteins related to classical detoxification strategies, including cytochrome P450 (N=6; Fig. 4A), GSTs (N=26; Fig. 4B), glycosyltransferases (N=13; Fig. 4C), xanthine dehydrogenase (N=1; Fig. 4D), and aldehyde oxidase (N=1; Fig. 4E) compared to the untreated control. These findings highlight possible temporal regulation of classical defense responses to *F. graminearum* in wheat prior to 21 dpi and demonstrate strain-specific defense activation routes used by the plant to protect from infection.

**Figure 3.**
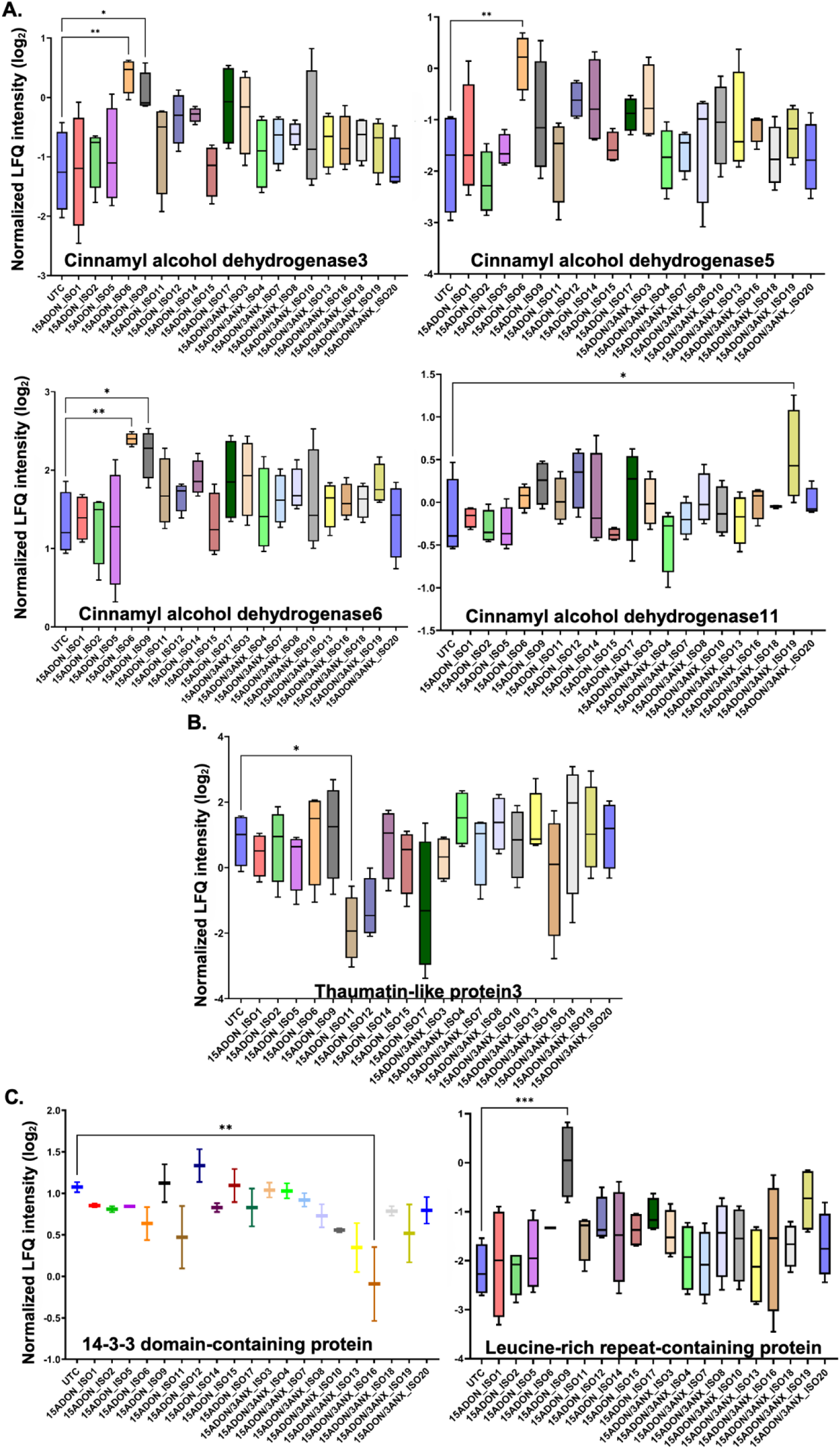
Variation in wheat defence□ and signalling□related protein abundance across *F. graminearum* strains producing 15ADON or 15ADON/3ANX. **A.** Proteins associated with lignin biosynthesis. **B.** Proteins involved in pathogen responsive defence pathways. **C.** Proteins linked to signal transduction. Bars represent the mean LFQ intensity for each condition, and error bars indicate the standard deviation calculated from four biological replicates. Differential protein abundance was assessed using one way ANOVA followed by Dunnett’s multiple comparison test (p < 0.05) relative to the untreated control.

**Figure 4.**
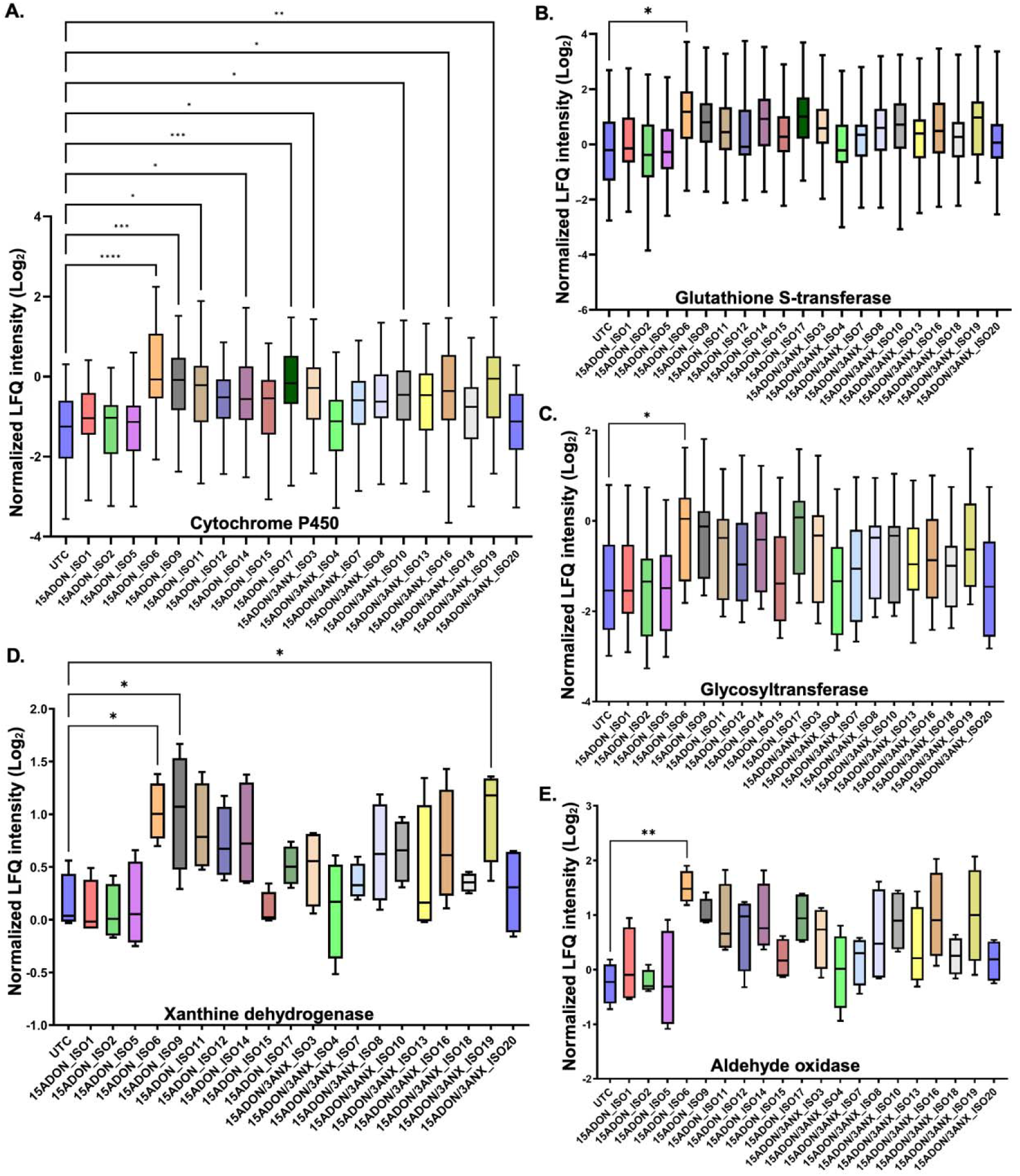
Variation in wheat detoxification□associated proteins across *Fusarium graminearum* strains producing 15ADON or 15ADON/3ANX. **A.** Cytochrome P450s, **B.** Glutathione S transferases, **C.** Glycosyltransferases, **D.** Xanthine dehydrogenase, and **E.** Aldehyde oxidase. Bars represent mean LFQ intensities, and error bars indicate standard deviation from four biological replicates per strain. These profiles highlight strain specific modulation of classical DON detoxification mechanisms. Protein abundance differences were evaluated using one way ANOVA followed by Dunnett’s multiple comparison test (p < 0.05) relative to the untreated control.

### Chemotype-exclusive fungal proteome reveals activation of 3ANX-specific pathways during wheat infection

To move beyond the host response and identify putative fungal proteins influenced by the presence and production of the 3ANX chemotype, we explored the data from the fungal perspective. For instance, a PCA of the common fungal proteome across chemotypes showed distinction across component 1 (86.46%) and component 2 (2.13%). However, the clusters were not associated with mycotoxin chemotype, suggesting other drivers (e.g., strain source, environmental factors) of fungal proteome remodelling (Fig. 5A). To examine replicate reproducibility, we assessed Pearson correlation values for each chemotype with 67.0% for 15ADON-treated samples and 76.3% for 15ADON/3ANX-treated samples (Fig. S3). As expected, these values are lower than the replicate reproducibility of the wheat proteome, given diverse strains are used for each chemotype, suggesting increased variability of fungal protein production and abundance across the strains and over the time course of infection. An observation of dynamic range for the fungal proteome highlighted similar detection for both 15ADON/3ANX- and 15ADON-producing strains, covering over two orders of magnitude in protein abundance (Fig. 5B).

**Figure 5.**
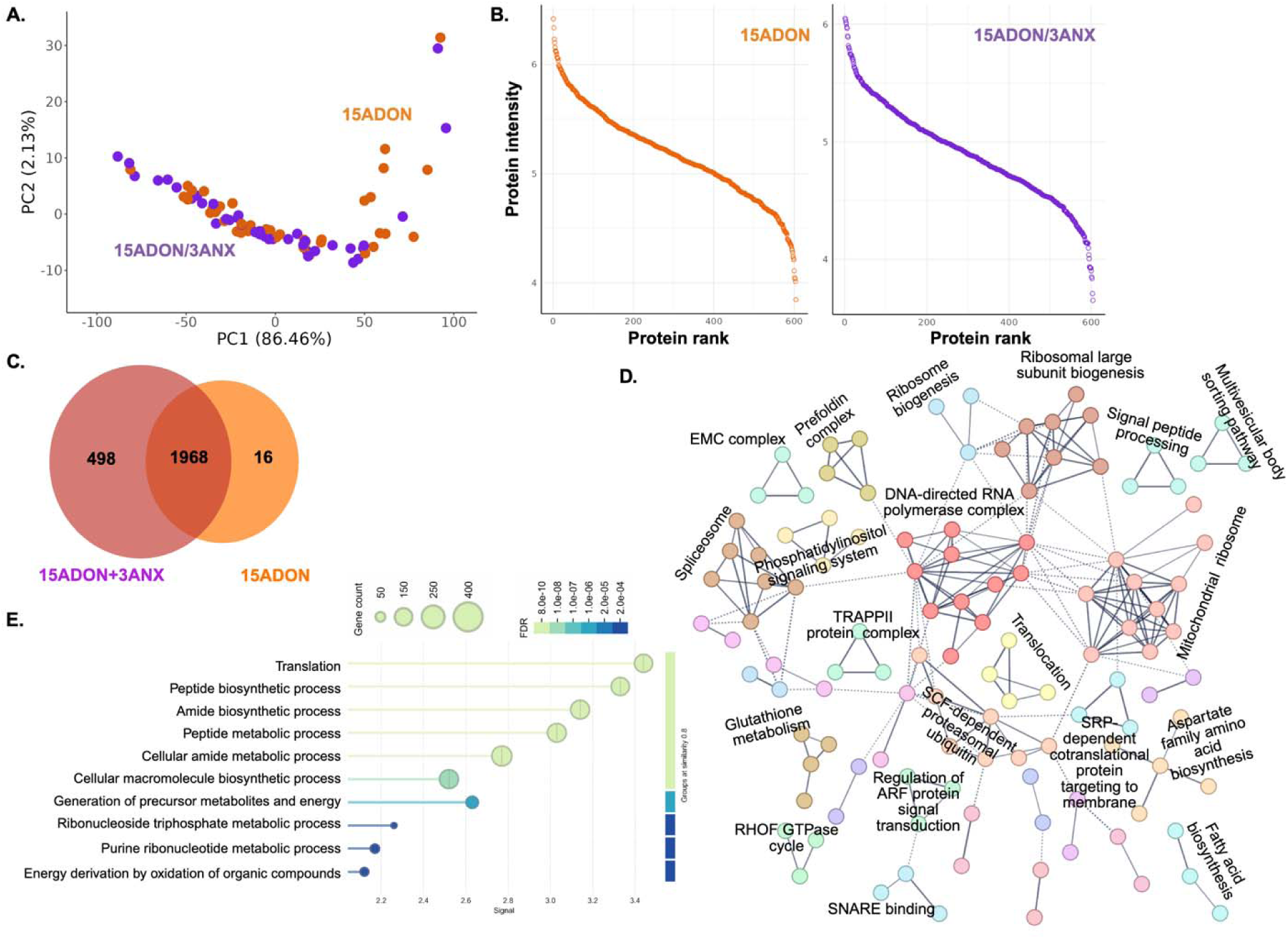
Chemotype-specific remodeling of the *Fusarium graminearum* proteome. **A.** Principal component analysis illustrating variation in fungal protein abundance across strain producing 15ADON or 15ADON/3ANX. **B.** Dynamic range plots showing the distribution of fungal protein intensities for each chemotype, demonstrating comparable detection depth across strains. **C.** Venn diagram depicting fungal proteins shared between chemotypes and those uniquely detected. **D.** STRING interaction network of proteins exclusive to the 15ADON/3ANX chemotype, clustered using the Markov Cluster Algorithm (MCL); 44 clusters defined with top 20 labeled. **E.** Gene Ontology Biological Process (GOBP) enrichment analysis of 15ADON/3ANX-exclusive proteins.

Next, we identified common proteins across fungal chemotypes (N = 1968), while observing 498 fungal proteins exclusively produced by the 15ADON/3ANX chemotype and 16 proteins exclusive to the 15ADON chemotype (Table S4; Fig. 5C). A detailed analysis of the 498 proteins exclusive to 15ADON/3ANX using STRING interaction network mapping and Markov Cluster Algorithm (MCL)-based clustering revealed 44 distinct clusters (Fig. 5D). These clusters prominently featured proteins involved in mitochondrial ribosome function, ribosomal large subunit biogenesis, and various metabolic pathways, including glutathione metabolism, which is associated with fungal oxidative stress resistance and pathogenicity (Park *et al*., 2024), as well as amino acid biosynthesis (e.g., aspartate family). Additional clusters highlighted roles in intracellular transport, translocation mechanisms, such as signal recognition particle (SRP)-dependent and co-translational targeting to membranes, and biosynthetic processes related to fatty acids and signal peptide processing. Complexes, such as the prefoldin, EMC (Endoplasmic Reticulum Membrane Complex), and TRAPPII were also notably represented, underscoring the multifaceted cellular roles of these proteins. Importantly, these latter categories have defined roles in cell wall integrity, fungal virulence, and suppression of plant immunity, respectively (Tahmaz *et al*., 2022; Ma *et al*., 2023; Liu *et al*., 2025). A complementary GOBP enrichment plot showcased enrichment across translation, biosynthetic and metabolic processes, as well as metabolite production and energy derivation (Fig. 5E). These findings indicate a chemotype-exclusive *F. graminearum* proteome produced in the presence of 3ANX and suggest protein interactions and pathways that are activated to protect from wheat defenses during host colonization.

### Chemotype-dependent remodeling of fungal metabolic and effector proteins during wheat infection

From the fungal perspective, we next asked whether virulence- and metabolism-associated proteins showed consistent trends when strains were grouped by mycotoxin chemotype rather than compared individually. Using the fungal proteome (604 proteins detected across all strains), we grouped samples into 15ADON- and 15ADON/3ANX-producing strains and compared protein abundance distributions between chemotypes. Here, protein abundance profiles did not differ significantly between chemotypes, demonstrating a consistent baseline of infection machinery (Fig. 6A); however, a subset of metabolic enzymes and a putative effector displayed significant chemotype-dependent differences. For example, cutinase (EC 3.1.1.74), a cell wall–degrading enzyme implicated in host surface penetration (Feng *et al*., 2005), showed significantly higher abundance in 15ADON strains compared to 15ADON/3ANX strains (Fig. 6B), suggesting that classical 15ADON producers may rely more heavily on cuticle degradation during infection. In contrast, enolase (EC 4.2.1.11), a key glycolytic enzyme (Gao *et al*., 2023), was significantly more abundant in 15ADON/3ANX strains (Fig.6C), indicating enhanced primary carbon metabolism and energy generation in emerging chemotypes. Additionally, strain specific analysis of enolase abundance revealed substantial variability across isolates, with 15ADON_ISO2 exhibiting the lowest enolase abundance and 15ADON/3ANX_ISO20 showing the highest levels among all strains (Figure S4A). Similarly, the effector CFEM1 (Common in Fungal Extracellular Membrane) (Chen *et al*., 2021) exhibited increased abundance in 15ADON/3ANX strains relative to 15ADON strains (Fig 6D), consistent with a shift toward effector-driven modulation of host immunity in the presence of 3ANX.

**Figure 6.**
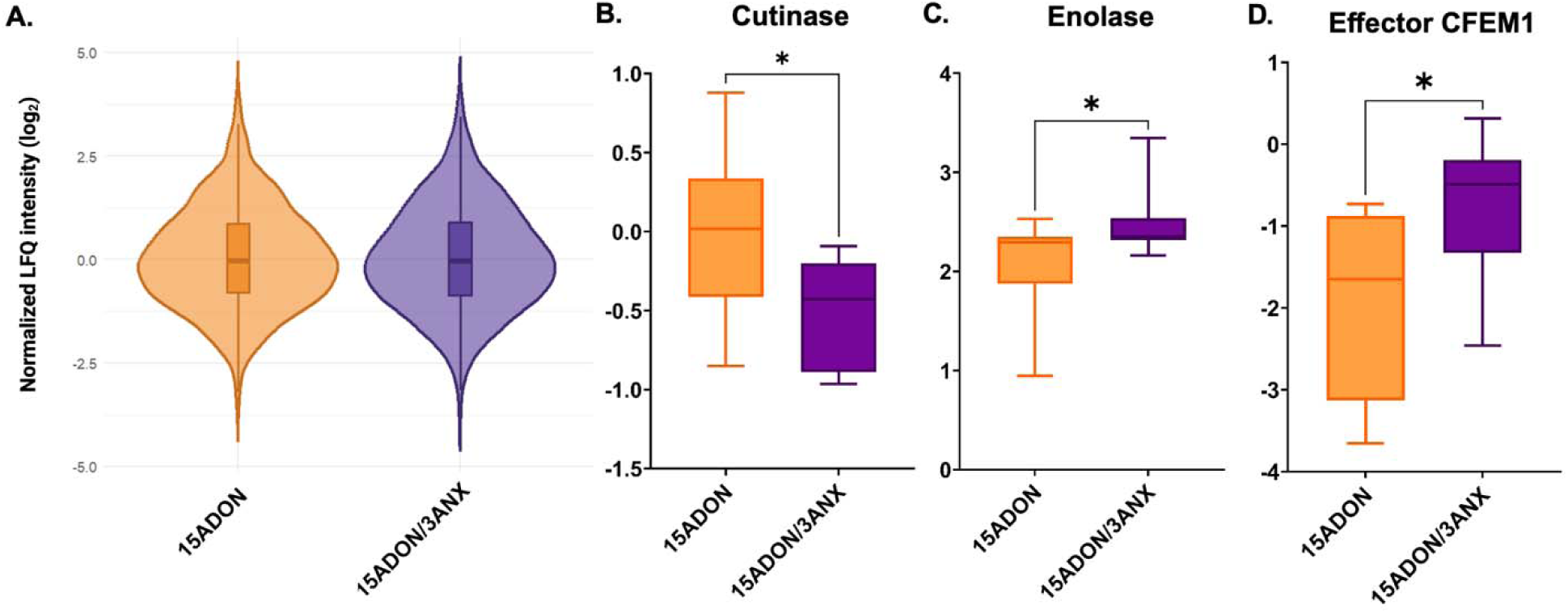
Chemotype-dependent differences in fungal protein abundance during wheat infection. Abundance of fungal proteins by: **A.** Global proteome intensities by chemotype, **B.** Cutinase (EC 3.1.1.74), **C.** Enolase (EC 4.2.1.11), and **D.** Effector CFEM1 in wheat heads infected with 15ADON or 15ADON/3ANX producing *F. graminearum* strains. Data are presented as boxplots showing the distribution of log transformed protein intensities for each chemotype group. Asterisks indicate significant differences between chemotypes (p < 0.05).

Next, given our previous observations of differential ergosterol production associated with the 3ANX chemotype (Ramezanpour *et al*., 2026), we also assessed five ergosterol associated proteins; however, none showed significant differences in abundance between chemotypes or among individual isolates (Figure S4B-F). This outcome is likely influenced by substantial biological variability inherent to field grown samples, the late sampling stage (21 dpi) when sterol biosynthesis may be less dynamic, and the lack of *in vitro* baseline measurements needed to distinguish true strain specific differences. Together, the 3ANX-exclusive fungal proteome suggests a model in which emerging 15ADON/3ANX strains reconfigure their virulence strategy by relying on metabolic flexibility and effector deployment to sustain infection and counteract host defences, reinforcing the concept of chemotype-specific fungal adaptation during wheat colonization.

### Protein interaction networks identify key regulators of detoxification in the wheat-Fusarium graminearum pathosystem

Given our observations of strain-specific proteome remodeling across both biological systems, we performed a network analysis to identify hub proteins and their associated interactors to further understand mechanisms driving the wheat-FHB interaction. Using 12 protein interaction networking mapping methods, 38 wheat proteins were identified with the highest number of interactions (Fig. 7A; Table S5), including 17 hub proteins belonging to various units and subunits of ribosomal proteins, indicating that the pathogen may disrupt host translation to facilitate infection. Conversely, these hubs may correlate with increased wheat ribosomal processes to mount effective defenses, given the dual role of ribosomal proteins in translation and stress signaling (Ramu *et al*., 2020; Reynoso, 2025). The differential abundance of hub proteins across all strains compared to the untreated control was assessed using ANOVA and three protein showed significant regulatory differences: DSK2a (DOMINANT SUPPRESSOR OF KAR 2a), eEF1γ2 (Elongation factor 1-gamma 2), and an uncharacterized protein (with UniProt accession A0A3B5XUE5) (Fig. 7B).

**Figure 7.**
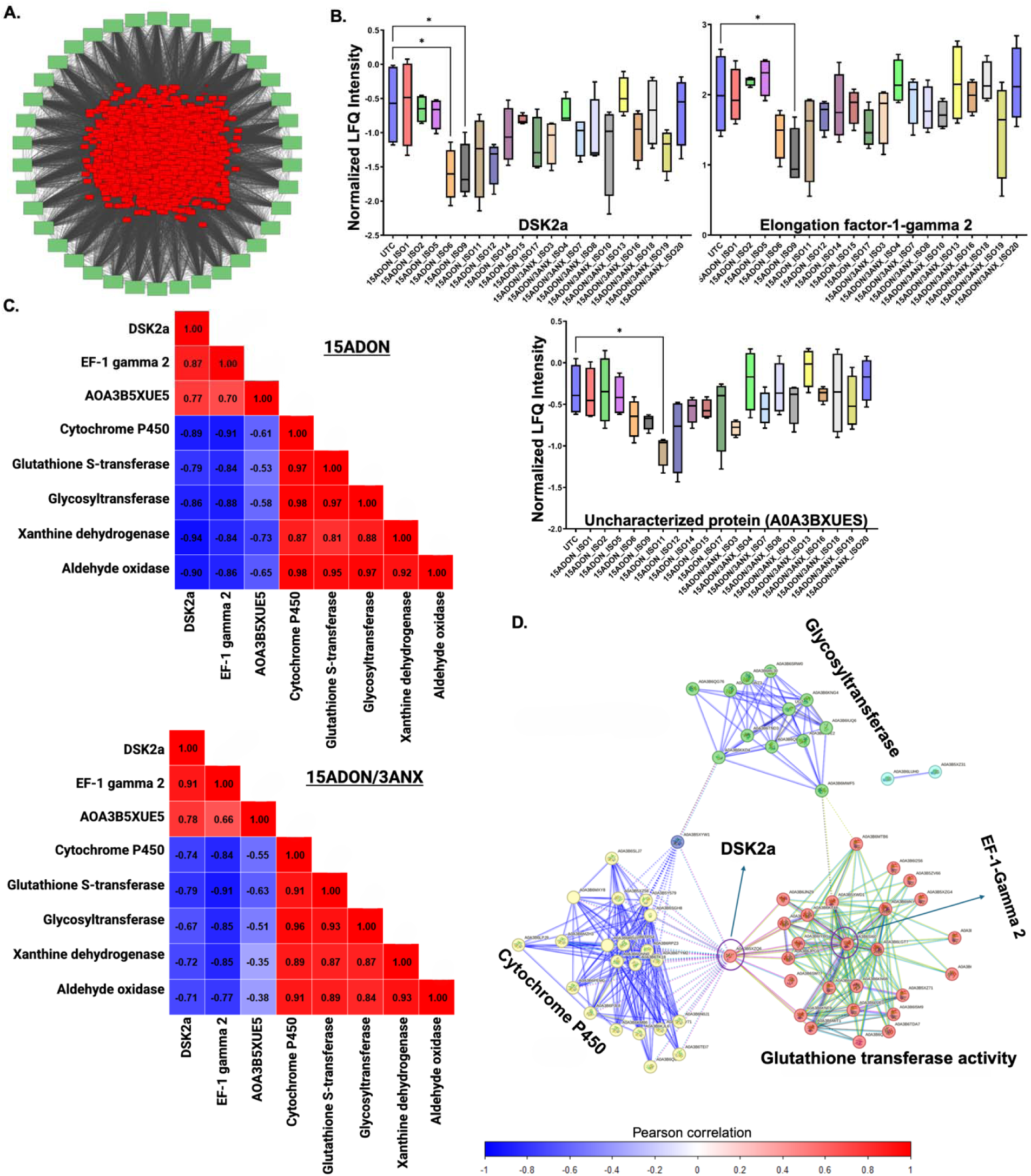
Integrated network, abundance, and correlation analyses reveal chemotype□dependent regulation of wheat hub proteins. **A.** Core wheat protein interaction network highlighting central regulatory architecture. Red nodes represent core proteins shared across treatments, while peripheral green nodes denote the 38 hub proteins identified using CytoHubba scoring applied to STRING based interaction networks. Edges reflect high confidence (0.7) predicted protein–protein interactions. **B.** Abundance profiles of selected hub proteins (DSK2a, elongation factor 1 gamma 2, and uncharacterized protein AOA3BXUES) across treatment groups. Box plots show mean LFQ intensities with standard deviation from four biological replicates. Statistical significance was evaluated using one way ANOVA followed by Dunnett’s post hoc test (p < 0.05). **C.** Pearson correlation heatmaps illustrating relationships among core proteins, hub regulators, and detoxification associated proteins under 15ADON and 15ADON/3ANX treatments. Correlation values range from negative (blue) to positive (red), demonstrating chemotype specific shifts in regulatory connectivity; values indicated within heatmap. **D.** STRING interaction network integrating hub proteins with detoxification related proteins, the uncharacterized hub protein A0A3B5XUE5 showed no detectable interactions within this detoxification-associated network. Functional clusters are color coded and annotated (e.g., glycosyltransferases, cytochrome P450s, glutathione transferases).

Next, we explored the relationship between the three hub proteins and five known mycotoxin detoxification proteins: cytochrome P450, glycosyltransferase, GST, aldehyde oxidase, and xanthin dehydrogenase. We observed a negative correlation between hub protein abundance and abundance of detoxification proteins, suggesting that the hub proteins may negatively regulate detoxification pathways in wheat-FHB interactions independent of the presence of 3ANX (Fig 7C). A tailored STRING network analysis of the hub proteins and detoxification proteins identified DSK2a to be connected by cytochrome P450 and GST clusters and eEF1γ2 within the centre of the GST network, whereas the uncharacterized hub protein A0A3B5XUE5 showed no detectable interactions with these detoxification-associated clusters (Fig. 7D). The inverse abundance patterns between the selected hub proteins and detoxification enzymes indicates a potential negative regulatory relationship, where activation of proteostasis and translational hubs may suppress canonical detoxification pathways, thereby reshaping wheat’s defense strategy during FHB infection.

### Chemotype-specific detoxification strategies in wheat: hub proteins suppress defense while 3ANX alters detoxification mechanisms

Given the observed regulatory divergence across wheat and pathogen proteomes in response to mycotoxin chemotypes and specific fungal strains, we aimed to investigate a connection across mycotoxin detoxification and accumulation and severity in wheat. We measured the D3G/DON ratio and disease severity for each fungal strain at the field collection time point (21 dpi). We observed comparable ratios of D3G/DON between wheat inoculated with 15ADON vs. 15ADON/3ANX, suggesting that any differences in the conversion of DON to D3G are strain-specific and not global (Fig. 8A). For disease severity, the 15ADON inoculated wheat showed a higher average value (60.0%) compared to 15ADON/3ANX (55.9%) but these levels were not significantly different (Fig. 8B). Intriguingly, variance in disease severity for 15ADON- vs. 15ADON/3ANX-inoculated wheat was 2.5-fold higher, indicating less consistent symptom formation in the absence of 3ANX.

**Figure 8.**
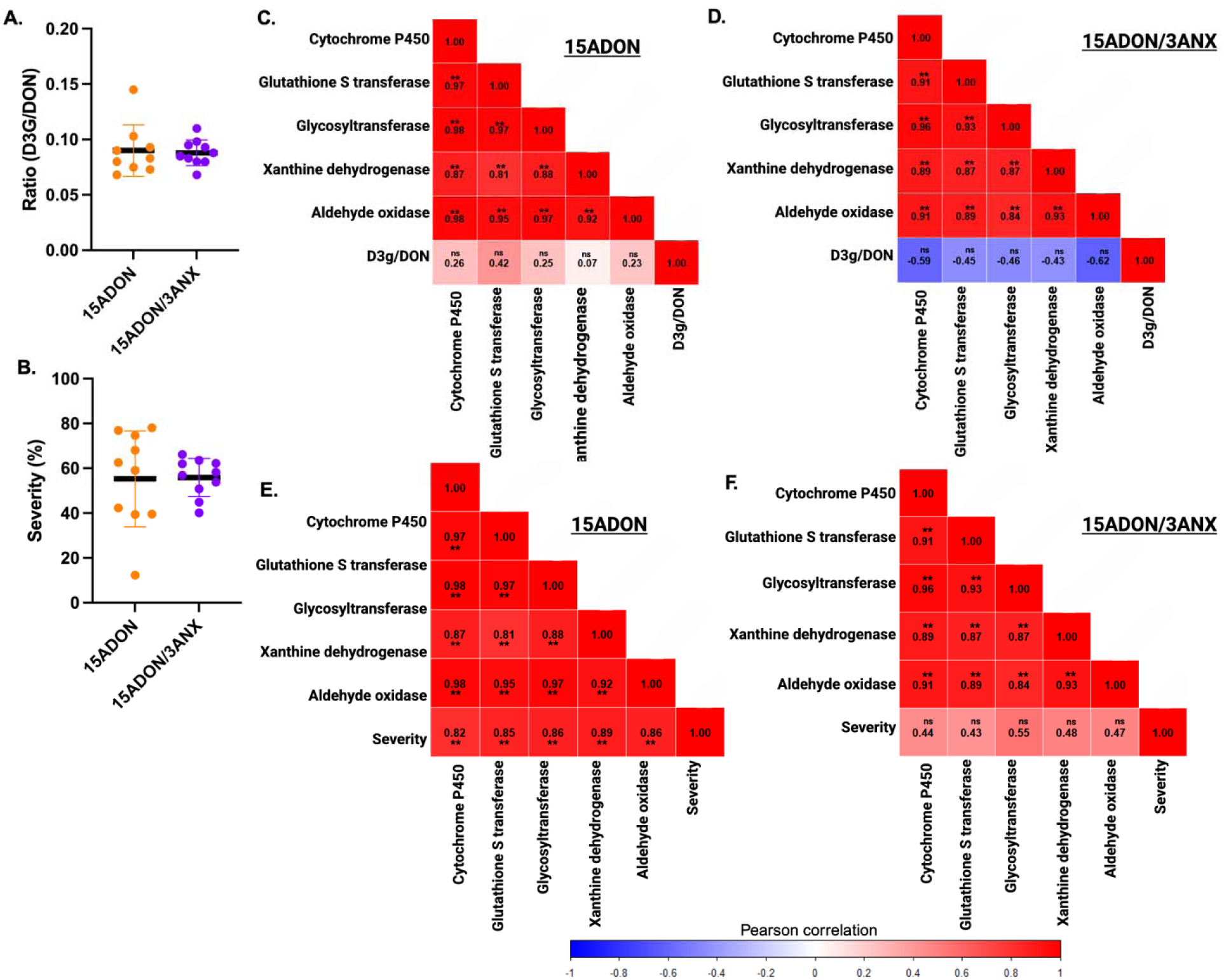
Chemotype□dependent correlation networks linking wheat detoxification proteins, DON conversion, and disease severity. **A.** Ratio of measured D3G/DON for wheat inoculated with 15ADON (orange) or 15ADON/3ANX (purple). **B.** Disease severity percentage for wheat inoculated with 15ADON (orange) or 15ADON/3ANX (purple). Average values (standard deviation) across fungal strains. Correlation matrices illustrate relationships among detoxification associated proteins and the D3G/DON ratio: **C.** 15ADON producing strains and **D.** 15ADON/3ANX producing strains, and the detoxification associated proteins. Correlation matrices illustrate relationships among detoxification associated proteins and disease severity: **E.** 15ADON producing strains and **F.** 15ADON/3ANX producing strains. Pearson correlation coefficients are shown, with significance indicated as p < 0.05 (*), p < 0.01 (**), and non significant (ns).

Next, to assess potential regulatory mechanisms driving mycotoxin accumulation and disease severity, we mapped protein abundance profiles of prioritized mycotoxin-associated proteins to these values. Wheat inoculated with 15ADON-producing strains showed a positive correlation with the D3G/DON value, indicating that classical, elevated detoxification activity aligned with increased conversion of DON to its less toxic, glycosylated form (D3G) (Fig. 8C). This conversion is a protective mechanism used by the plant to detoxify the mycotoxin and limit damage to grains (Simsek *et al*., 2012; Ovando-Martínez *et al*., 2013; Nagl *et al*., 2014; Tucker *et al*., 2019; Guo *et al*., 2020). Conversely, wheat inoculated with 15ADON/3ANX-producing strains displayed the opposite pattern with abundance of detoxification proteins negatively correlated with the D3G/DON value (Fig. 8D). These data indicate that in the presence of 3ANX, detoxification pathways may shift away from classical DON detoxification systems toward alternative mechanisms for the conversion of DON into D3G. Additionally, we examined disease severity and observed that wheat inoculated with 15ADON-producing strains showed a strong positive correlation (>0.82; p<0.01) with detoxification protein abundance (Fig. 8E); however, inoculation with 15ADON/3ANX-producing strains did not indicate a correlation (0.43 – 0.55) between disease and detoxification (Fig. 8F). These data further support our hypothesis that 3ANX evades or suppresses classical defense mechanisms, such as we observed above through inverse regulation between hub proteins and key detoxification proteins, disrupting the relationship between DON metabolism, detoxification approaches, and disease severity.

## Discussion

Our study sheds new light on the molecular interactions between *T. aestivum* and *F. graminearum* under the selection pressure of different trichothecene chemotypes, particularly the developing 3ANX variation in combination with 15ADON. By profiling host and pathogen proteins from field-grown wheat spikes three weeks post inoculation, we record a late-stage interaction (21 dpi) in which both species engaged in significant physiological and metabolic adaptations. This temporal context is critical as at this time, the proteome (transcriptome and genome) is more likely to reflect protracted defence responses, structural strengthening, and long-term metabolic reprogramming, rather than the quick, transitory signalling processes seen in early infection (Seifi *et al*., 2023; Buchanan *et al*., 2025; Walker *et al*., 2025). Pathogen detection, oxidative bursts, and activation of primary defensive cascades often drive early-stage responses (hours to days after inoculation). In contrast, the late responses described herein may be unique to the chronic infection phase, such as the detoxification and accumulation of mycotoxins as disease severity increases.

We expand upon our previous findings (Ramezanpour *et al*., 2026) to focus on strain-specific responses, hub protein dynamics, and the relationship among chemotype, mycotoxin conversion, and disease severity. By updating our approach to leverage DIA-based high-resolution MS, we improved the depth of proteome coverage across host and pathogen from traditional DDA-based approaches by 1.8-fold. Comparison to our earlier work reveals both conserved and newly emerging features of wheat–*Fusarium* interactions. For instance, we confirmed the strong activation of oxidative and structural defense pathways, including peroxidases, enolase, and cell-wall-modifying enzymes, alongside the characteristic suppression of photosynthesis. However, enhanced proteome depth covered across phenotyping and correlation analyses provide a new study framework that uncovers additional chemotype-specific regulatory patterns. While our previous study (Ramezanpour *et al*., 2026) suggested reduced GST-mediated detoxification in specific 15ADON/3ANX strains, this current study demonstrates that this chemotype fundamentally disrupts the expected players in mycotoxin detoxification, accumulation, and disease severity. Specifically, we found that detoxification proteins correlate positively with disease severity in 15ADON infections, but this association is lost in 15ADON/3ANX infections. Furthermore, the identification of hub proteins, such as DSK2a, acting as a connector between regulatory subnetworks, and the observation of opposing trends in D3G/DON ratios provide new mechanistic insight into how 3ANX may suppress classical DON detoxification pathways. From the fungal perspective, while both studies identified 3ANX-exclusive proteomes enriched in virulence-associated pathways, the current dataset expands this to include mitochondrial ribosome components and ER–mitochondrion interface complexes. Together, these findings demonstrate that while core wheat defenses remain conserved, emerging 3ANX chemotypes uniquely reshape both host detoxification networks and fungal metabolic processes in ways not captured in prior investigations.

The enrichment of glutathione metabolism and ABC transporter activity across both chemotypes demonstrates the importance of oxidative stress reduction and mycotoxin detoxification in wheat-FHB defence. However, 3ANX-containing infections specifically stimulated chitinases, defensins, heat shock proteins (HSP26), and glycosyltransferases, indicating a shift toward structural barrier reinforcement, protease inhibition, and glycosylation-mediated mycotoxin detoxification (Park & Seo, 2015; El Mounadi *et al*., 2016; Shah *et al*., 2017; Xing *et al*., 2018). Our findings lend evidence in the idea that resistance to *F. graminearum* is achieved by coordinated modulation of defence proteins and stress-related pathways. For instance, previous integrative proteomics demonstrated that chloroplast-associated functions and early signalling events constitute a core responsive network across wheat genotypes, highlighting the conserved nature of these defence mechanisms (Fabre *et al*., 2021). The abundance of peroxisomal proteins in 3ANX treatments suggests roles in lipid-derived signalling and reactive oxygen species metabolism, which are compatible with peroxisomal involvement in jasmonate biosynthesis and defence (del Río & López-Huertas, 2016; Kao *et al*., 2018). These patterns could represent sustained or secondary defence responses that evolve over disease invasion. Additionally, previous metabolomics studies in the FHB-resistant wheat cultivar Sumai-3 revealed similar defence signatures, including enhanced phenylpropanoid metabolism, cell wall fortification, and mycotoxin detoxification (Gunnaiah & Kushalappa, 2014), implying that these biochemical pathways are conserved across multiple layers of the plant’s defence system.

The 15ADON/3ANX-exclusive fungal proteome was enriched for mitochondrial ribosome components, mitochondrial interface complexes, TRAPPII vesicle trafficking machinery, and prefoldin chaperones, all of which are involved in protein homeostasis, secretion, and virulence. The predominance of glutathione metabolism and amino acid biosynthesis pathways suggests that 3ANX-producing strains improve redox buffering and metabolic flexibility to tolerate host-derived oxidative stress. The existence of cell wall integrity-associated complexes suggests that these chemotypes may enhance structural resilience during host colonization, possibly counteracting plant cell wall-degrading enzyme inhibitors. Our chemotype-level analysis of the fungal proteome further supports the concept that emerging 3ANX-producing strains deploy distinct virulence strategies during wheat infection. When fungal strains were grouped by chemotype, rather than compared individually, several consistent patterns emerged. Classical 15ADON strains exhibited higher abundance of cutinase, a key enzyme involved in host surface penetration, suggesting a greater reliance on physical degradation of host barriers. In contrast, 15ADON/3ANX strains showed significantly elevated abundance of enolase and the effector CFEM1, indicating a shift toward enhanced primary metabolism and effector-mediated modulation of host immunity. For enolase, the wide dynamic range suggests that individual isolates differ markedly in their glycolytic activity during wheat infection, potentially reflecting isolate specific metabolic strategies or growth rates. This trend was also consistent with our previous field study, where reduced production of enolase was observed in 15ADON_ISO2, confirming that ISO2 repeatedly exhibits the lowest enolase abundance (Ramezanpour *et al*., 2026). Fungal effectors, such as the observed CFEM1, have diverse roles in modulating infection through suppressing immune signaling, modulating oxidative stress responses, and altering host cell wall integrity(Chen et al., 2021). These molecules act in both the apoplast and cytoplasm to reprogram host physiology, thereby facilitating tissue colonization and mycotoxin accumulation (Rauwane *et al*., 2020). These chemotype-dependent differences align with the broader 3ANX-exclusive fungal proteome, which was enriched for mitochondrial ribosome components, secretory pathway complexes, and metabolic remodeling. Together, these findings suggest that 3ANX-producing strains prioritize metabolic flexibility and immune interference over classical cell-wall–degrading strategies, reflecting an adaptive reprogramming of fungal virulence in response to host defenses.

Network analysis revealed that two wheat hub proteins, DSK2a and EF1γ2, were inversely correlated with detoxification enzymes (i.e., cytochrome P450s, GSTs, glycosyltransferases, aldehyde oxidase, xanthine dehydrogenase) regardless of chemotype. DSK2, a conserved ubiquitin-binding receptor, mediates protein degradation in yeast, animals, and plants (Nolan *et al*., 2017); the clustering with cytochrome P450s and GSTs suggests that ubiquitin mediated proteostasis and protein quality control mechanisms may intersect with mycotoxin detoxification processes. These findings are consistent with evidence that proteostasis regulators modulate plant defense signaling and metabolic reprogramming under pathogen attack (Trujillo & Shirasu, 2010; Marino *et al*., 2012). Likewise, the placement of EF1γ2 within the GST associated cluster supports its emerging role in redox regulation and translational control during stress (Koonin *et al*., 1994). Given that eEF1γ contains a GST-like domain and DSK2 is a ubiquitin-binding receptor involved in selective protein degradation, these hubs may act as negative regulators of detoxification capacity, either by modulating enzyme turnover or by diverting cellular resources toward other stress responses. In contrast, the third hub protein, A0A3B5XUE5, remained unconnected to detoxification associated clusters, suggesting that its centrality within the broader network reflects functions unrelated to classical detoxification pathways and may represent an unexplored regulatory node activated during chemotype specific stress. Manipulating hub–detoxifier interactions could therefore enhance chemotype-specific resistance.

## Conclusions and future directions

Overall, our integrated comparative host–pathogen proteome map demonstrates that FHB resistance is a multifaceted trait, influenced by chemotype- and isolate-dependent proteomic interactions that shift dynamically throughout the infection process. The emergence of 3ANX producing chemotypes, with their distinct host and pathogen signatures, suggests that breeding programs must incorporate screening against multiple chemotypes and consider both early and late defence phases to capture durable resistance. Similar proteomics-based approaches have pinpointed defence-associated proteins, providing direct molecular targets for selection. Our observation of divergence of correlation between known mycotoxin detoxification-associated protein abundances and the ratio of D3G/DON in the presence of 3ANX is an important finding as it moves the work beyond descriptive regulation and into quantifiable biochemical differences that directly impact food safety and quality. Generating information about the suppression of classical defense responses in wheat upon challenge with 3ANX-producing fungal strains, along with identification of alternative defense responses, increases our knowledge of fungal virulence, along with putative biomarkers for selected breeding toward 3ANX-resistant cultivars (e.g., chitinases, defensins, glycosyltransferases), and defenses against conserved virulence factors, such as chitin synthase 3B and thioredoxin reductase, produced by the pathogen.

## Author Contributions

D.H. & J.G.-M. conceptualized and designed the study. S.R., N.A., & A.D. performed experiments, including field inoculations, sample collections, and mycotoxin analysis, protein extractions and sample preparations, proteomics sample preparation and mass spectrometry measurements. S.C. supported mass spectrometry experiments and bioinformatics analysis. S.R., J.A.M. & J.G.-M. analyzed data and generated figures and tables. S.R. & J.G.-M. wrote the first manuscript draft. All authors contributed to manuscript preparation and have read and approved the submitted manuscript.

## Acknowledgements

The authors thank Dr. Art W. Schaafsma for his valuable contributions and vision for improvements to Fusarium head blight management and critical assessment of mycotoxin production and contamination. We also thank members of the Geddes-McAlister lab for helpful discussions and constructive comments on the study. The authors thank Victor Limay-Rios for technical field assistance and Angela Connolly for mass spectrometry support. This work was facilitated by access to Sydney Mass Spectrometry, a Core Facility of The University of Sydney.

## Data Availability

The proteomics datasets are publicly available through PRIDE Proteomics Exchange:

For review purposes, please use:

Project accession: PXD073502

Token: N7myEcwuyxwd

Alternatively: reviewer can access the dataset by logging in to the PRIDE website using the following account details:

Username

Password: Qo3fhawyxUxv

## Supplemental Files

Table S1: Experimental design and *F. graminearum* strains.

Table S2: Wheat proteins exclusively detected upon 15ADON/3ANX chemotype treatment.

Table S3: Prioritized core wheat proteins with defined roles in pathogen defense and mycotoxin detoxification.

Table S4: Fungal proteins exclusively detected upon 15ADON/3ANX chemotype treatment. Table S5: List of core hub wheat proteins.

Figure S1: Wheat proteome column correlation by hierarchical clustering of Pearson correlation across samples.

Figure S2: Stable production of defense-related proteins in wheat in response to fungal chemotypes.

Figure S3: Fungal proteome column correlation by hierarchical clustering of Pearson correlation across inoculated samples.

Figure S4: Strain-specific abundance of enolase and ergosterol-associated proteins in *Fusarium graminearum*.

