## Supplementary material for "Recently emerged *Fusarium* chemotypes reprogram wheat defence and detoxification networks during Fusarium head blight development": Supp Fig

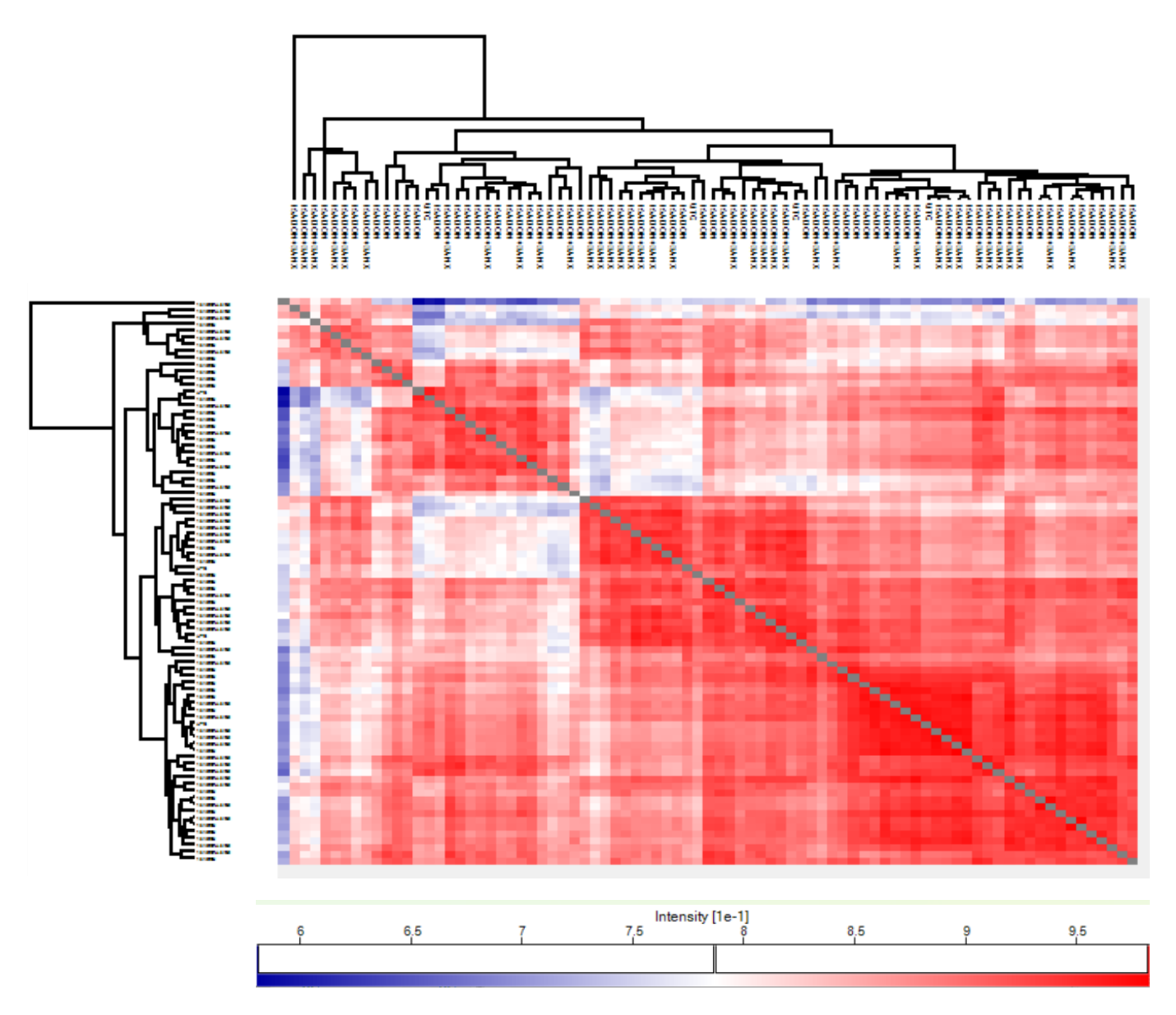

**Figure S1. Reproducibility of wheat proteome profiles across treatments based on hierarchical clustering of Pearson correlation values.** Correlation matrices illustrate sample‑to‑sample similarity for untreated wheat and for plants inoculated with 15ADON or 15ADON/3ANX chemotypes. Replicate reproducibility values were calculated as follows: control = 86.8%, 15ADON = 88.5%, and 15ADON/3ANX = 86.3%. Each chemotype includes four biological replicates composed of pooled wheat heads; the control group consists of four individual heads.

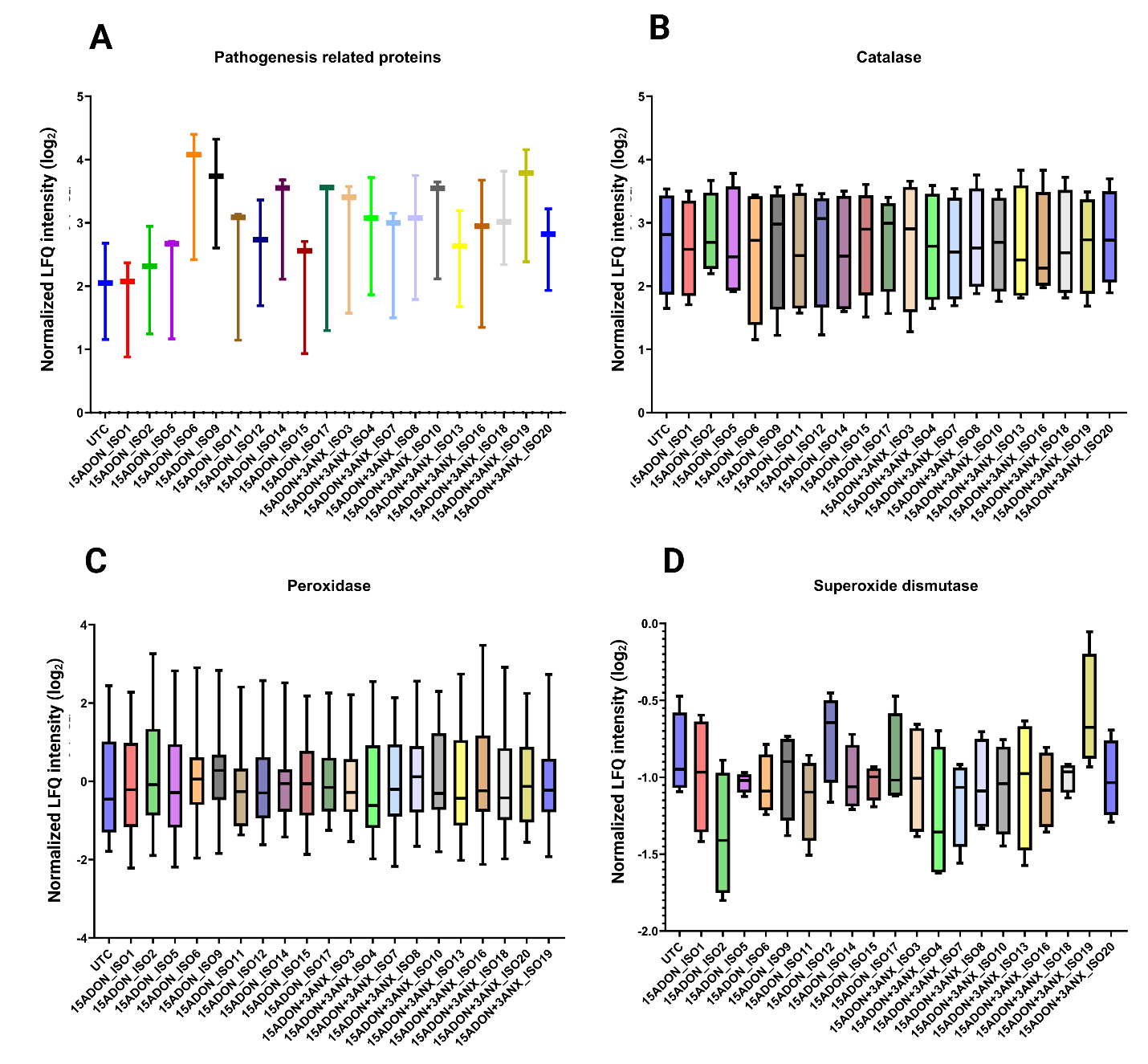

**Figure S2. Stable accumulation of selected wheat defense‑related proteins following infection with different *Fusarium* chemotypes. A.** Abundance profiles of pathogenesis‑related proteins (N = 3). **B.** Catalase abundance across treatments (N = 4). **C.** Peroxidase family proteins (N = 19).

**D.** Superoxide dismutase (N = 1). Data represent mean LFQ intensities with standard deviation across four biological replicates per treatment. Differential protein abundance was assessed using one‑way ANOVA followed by Dunnett’s multiple comparison test (p < 0.05) relative to the untreated control.

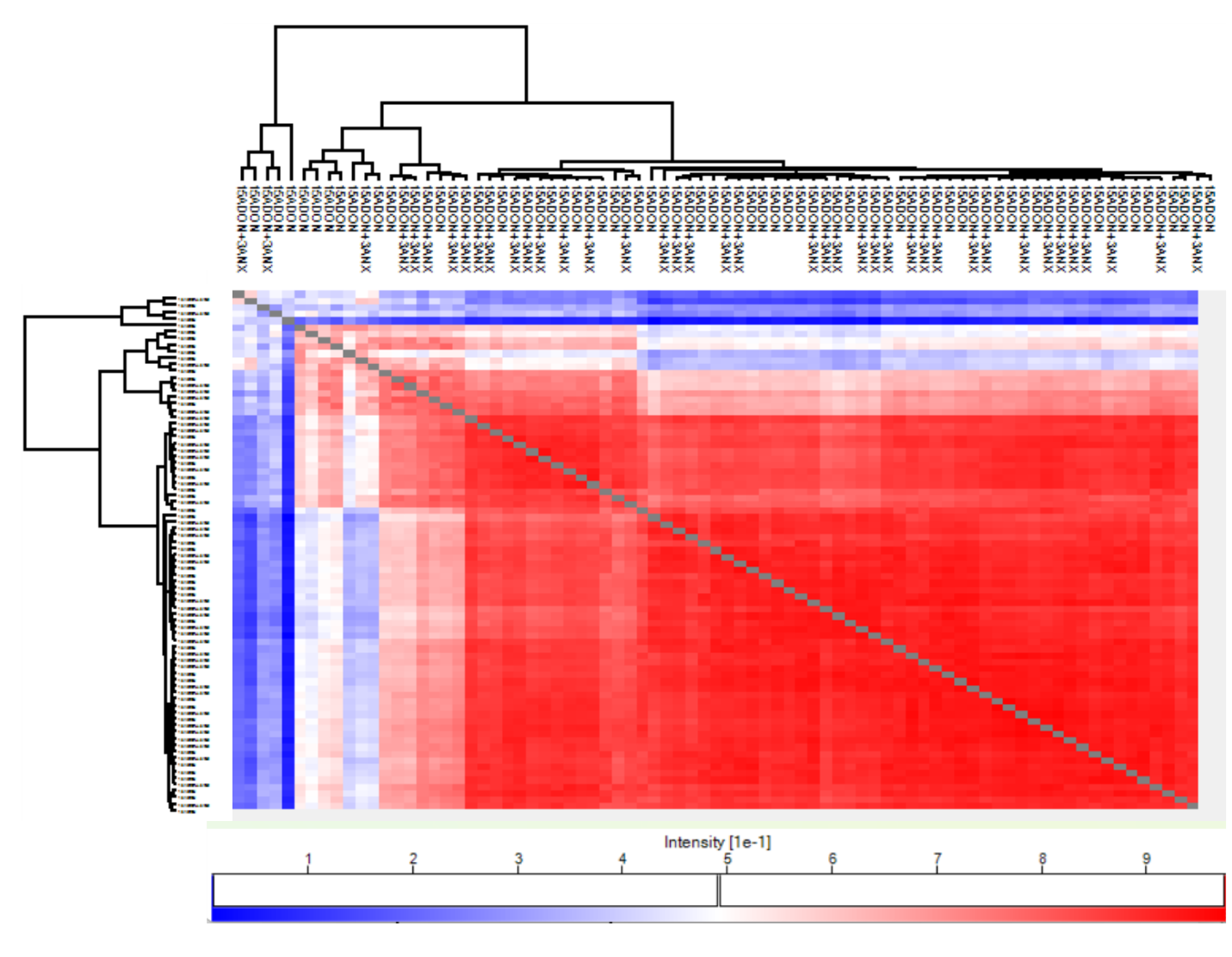

**Figure S3. Consistency of fungal proteome measurements across inoculated samples assessed by hierarchical clustering of Pearson correlation coefficients.** Correlation patterns and reproducibility metrics are shown for fungal proteins detected in wheat infected with 15ADON or 15ADON/3ANX isolates. Replicate reproducibility was 67.0% for 15ADON and 76.3% for 15ADON/3ANX. Each chemotype includes four biological replicates derived from pooled wheat heads.

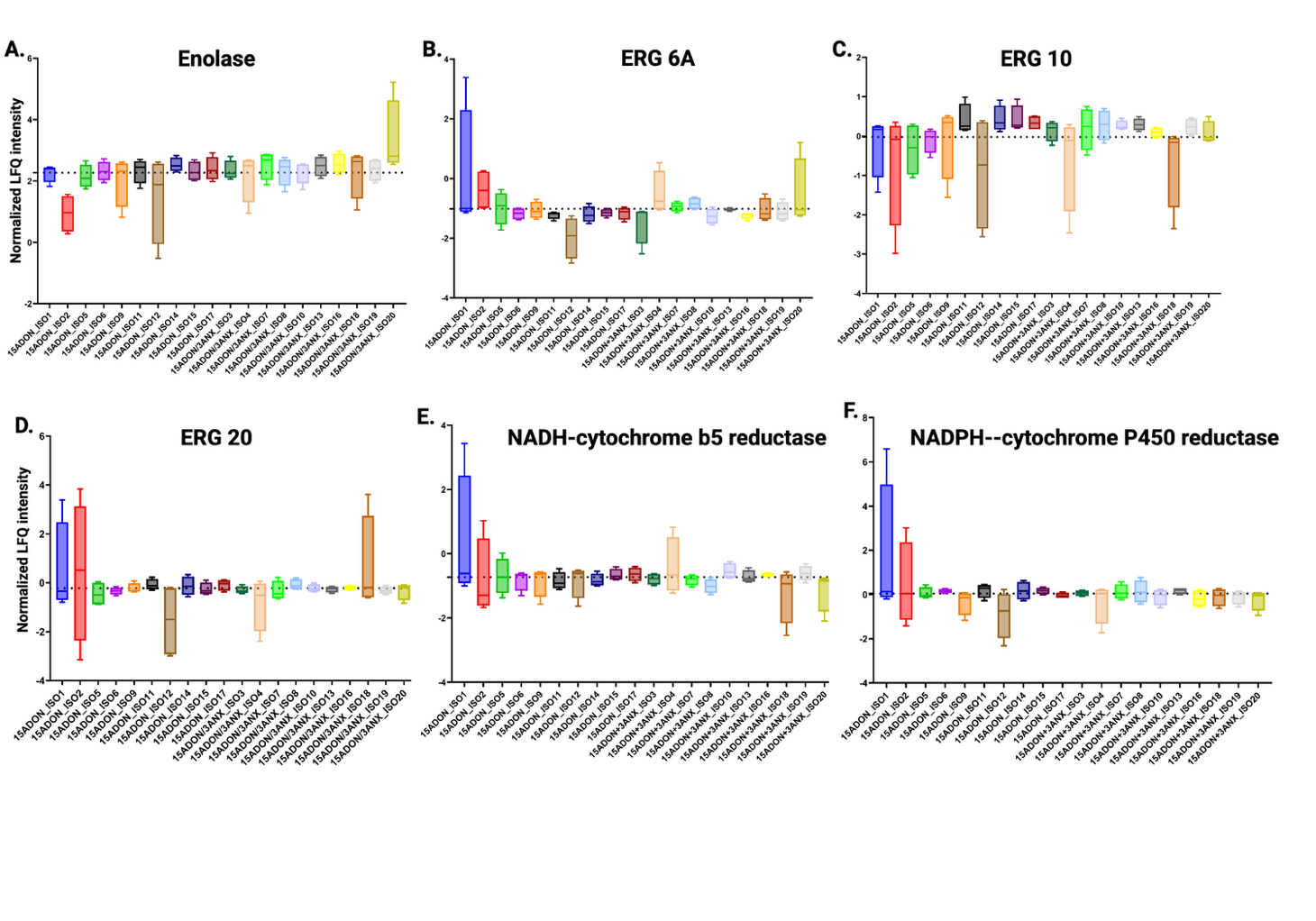

**Figure S4. Strain‑specific abundance of enolase and ergosterol‑associated proteins in *Fusarium graminearum* during wheat infection.** Normalized LFQ intensities for A) enolase and B-F) five ergosterol‑associated proteins (ERG6A, ERG10, ERG20, NADH–cytochrome b5 reductase, and NADPH–cytochrome P450 reductase). Each box plot represents protein abundance for individual isolates grouped by chemotype (15ADON or 15ADON/3ANX). LFQ values were log2‑transformed prior to visualization. The dashed line represents the mean LFQ intensity across all strains. Mean ± SD is shown for four biological replicates per condition.
