## Supplementary material for "Recently emerged *Fusarium* chemotypes reprogram wheat defence and detoxification networks during Fusarium head blight development": Supp Tables

**Table S1: Overview of *Fusarium graminearum* Chemotypes, Strains, and Inoculation Structure Used in the 2023 Field Experiment.**

| **Chemotype** | **Inoculum** | **Strain** | **DAOMC identifier** | **Host*** | **Source** | **Sample ID** |
| --- | --- | --- | --- | --- | --- | --- |
| 15ADON | Mycelium | 2 | 177406 |  | AAFC | 6 |
|  |  |  |  |  |  | 7 |
|  |  |  |  |  |  | 8 |
|  |  |  |  |  |  | 9 |
|  |  | 5 | 178149 |  | AAFC | 17 |
|  |  |  |  |  |  | 18 |
|  |  |  |  |  |  | 19 |
|  |  |  |  |  |  | 20 |
|  |  | 6 | 180376 |  | AAFC | 21 |
|  |  |  |  |  |  | 22 |
|  |  |  |  |  |  | 23 |
|  |  |  |  |  |  | 24 |
|  |  | 9 | 180379 |  | AAFC | 33 |
|  |  |  |  |  |  | 34 |
|  |  |  |  |  |  | 35 |
|  |  |  |  |  |  | 36 |
|  |  | 11 | 180377 |  | AAFC | 41 |
|  |  |  |  |  |  | 42 |
|  |  |  |  |  |  | 43 |
|  |  |  |  |  |  | 44 |
|  |  | 14 | 180378 |  | AAFC | 51 |
|  |  |  |  |  |  | 52 |
|  |  |  |  |  |  | 53 |
|  |  |  |  |  |  | 54 |
|  |  | 15 | 170785 |  | AAFC | 55 |
|  |  |  |  |  |  | 56 |
|  |  |  |  |  |  | 57 |
|  |  |  |  |  |  | 58 |
|  | Spore | 1 | 313121 | Stalk | RT | 1 |
|  |  |  |  |  |  | 3 |
|  |  |  |  |  |  | 4 |
|  |  |  |  |  |  | 5 |
|  |  | 12 | 252442 | Corn | RT | 45 |
|  |  |  |  |  |  | 46 |
|  |  |  |  |  |  | 47 |
|  |  |  |  |  |  | 84 |
|  |  | 17 | 252437 | Wheat | RT | 63 |
|  |  |  |  |  |  | 64 |
|  |  |  |  |  |  | 65 |
|  |  |  |  |  |  | 66 |
| 15ADON,3ANX | Spore | 3 | 252446 | Stalk | RT | 10 |
|  |  |  |  |  |  | 11 |
|  |  |  |  |  |  | 12 |
|  |  |  |  |  |  | 13 |
|  |  | 4 | 180377 | Stalk | RT | 14 |
|  |  |  |  |  |  | 15 |
|  |  |  |  |  |  | 16 |
|  |  |  |  |  |  | 82 |
|  |  | 7 | 252440 | Stalk | RT | 25 |
|  |  |  |  |  |  | 26 |
|  |  |  |  |  |  | 27 |
|  |  |  |  |  |  | 28 |
|  |  | 8 | 252441 | Wheat | RT | 29 |
|  |  |  |  |  |  | 30 |
|  |  |  |  |  |  | 31 |
|  |  |  |  |  |  | 32 |
|  |  | 10 | 252445 | Corn | RT | 37 |
|  |  |  |  |  |  | 38 |
|  |  |  |  |  |  | 39 |
|  |  |  |  |  |  | 40 |
|  |  | 13 | 252438 | Stalk | RT | 48 |
|  |  |  |  |  |  | 49 |
|  |  |  |  |  |  | 50 |
|  |  |  |  |  |  | 83 |
|  |  | 16 | 252439 | Corn | RT | 59 |
|  |  |  |  |  |  | 60 |
|  |  |  |  |  |  | 61 |
|  |  |  |  |  |  | 62 |
|  |  | 18 | N/A | Stalk | RT | 67 |
|  |  |  |  |  |  | 68 |
|  |  |  |  |  |  | 69 |
|  |  |  |  |  |  | 84 |
|  |  | 19 | 252443 | Stalk | RT | 70 |
|  |  |  |  |  |  | 71 |
|  |  |  |  |  |  | 72 |
|  |  |  |  |  |  | 73 |
|  |  | 20 | 252444 | Wheat | RT | 74 |
|  |  |  |  |  |  | 75 |
|  |  |  |  |  |  | 76 |
|  |  |  |  |  |  | 77 |
| Untreated |  |  |  |  |  | 78 |
|  |  |  |  |  |  | 79 |
|  |  |  |  |  |  | 80 |
|  |  |  |  |  |  | 81 |

DAOMC = Canadian Collection of Fungal Cultures

AAFC = Agriculture and Agri-Food Canada, Ottawa

RT = University of Guelph, Ridgetown campus

*Host = original source of pathogen

**Table S2: Chemotype‑Specific Wheat Proteins Exclusively Identified in 15ADON/3ANX‑Inoculated Samples**

| **Protein ID*** | **Protein name** | **Keyword/**  **Function^#^** | **LFQ Intensity (log2)** |
| --- | --- | --- | --- |
| A0A077RFH6 | (bread wheat) hypothetical protein | Hormone | 17.00 |
| A0A3B6RA92 | AAA+ ATPase domain-containing protein | Membrane | 12.21 |
| A0A3B6KIJ7 | AB hydrolase-1 domain-containing protein | Membrane | 12.83 |
| A0A3B6QCU8 | Actin-related protein 4 | - | 14.26 |
| A0A3B6KQP6 | Aldehyde oxidase | 2Fe-2S | 12.59 |
| A0A3B6SFP5 | aldehyde oxygenase (deformylating) (EC 4.1.99.5) | Endoplasmic reticulum | 13.75 |
| P01544 | Alpha-1-purothionin (Purothionin A-II) | 3D-structure | 20.80 |
| A0A3B6HRT6 | Apyrase | ATP-binding | 13.23 |
| A0A3B6R8E0 | Aromatic-L-amino-acid decarboxylase | Decarboxylase | 15.63 |
| A0A3B6RNN9 | ATP citrate synthase (EC 2.3.3.8) | Cytoplasm | 15.60 |
| A0A3B6NKF2 | BHLH domain-containing protein | - | 14.50 |
| A0A3B6KHH5 | Bifunctional inhibitor/plant lipid transfer protein | - | 15.76 |
| A0A3B6UA86 | Bifunctional inhibitor/plant lipid transfer protein | Lipid-binding | 18.15 |
| A0A3B6KL99 | BIG2 domain-containing protein | Glycoprotein | 13.64 |
| A0A3B5Y1X0 | Bowman-Birk serine protease inhibitors family domain-containing protein | - | 15.26 |
| A0A077S655 | BSD domain-containing protein | - | 13.60 |
| A0A3B6RBS3 | BZIP domain-containing protein | Abscisic acid signaling pathway | 13.74 |
| A0A3B6KS51 | C2 domain-containing protein | Membrane | 13.47 |
| A0A3B6KCN7 | C3H1-type domain-containing protein | DNA-binding | 13.10 |
| A0A3B6RHE7 | CAF1B/HIR1 beta-propeller domain-containing protein | Chromatin regulator | 14.24 |
| A0A3B6RBW2 | Carbohydrate kinase PfkB domain-containing protein | Kinase | 16.89 |
| A0A3B6HZH4 | Carbohydrate kinase PfkB domain-containing protein | Kinase | 15.00 |
| A0A3B6NNJ9 | Cathepsin propeptide inhibitor domain-containing protein | - | 15.54 |
| A0A3B6NNK3 | CBS domain-containing protein | CBS domain | 14.35 |
| A0A0A7LWQ8 | CCAAT-binding transcription factor A | - | 15.45 |
| A0A077RRA4 | Cell division cycle protein 123 | - | 12.96 |
| A0A3B5XZH6 | Cellulose synthase-like protein D3 | Cell wall biogenesis/degradation | 13.16 |
| W5QKZ0 | Chalcone-flavonone isomerase family protein | Flavonoid biosynthesis | 15.37 |
| A0A3B6SQ43 | chitinase (EC 3.2.1.14) | Carbohydrate metabolism | 16.49 |
| A0A3B6RDI5 | Chloride channel protein | CBS domain | 14.56 |
| Q9SBB7 | Chloroplast small heat shock protein | - | 15.66 |
| A0A3B6KGQ1 | CID domain-containing protein | Coiled coil | 13.27 |
| A0A3B6HXW1 | CID domain-containing protein | mRNA processing | 13.41 |
| A0A3B6MQ28 | Citrate transporter-like domain-containing protein | Antiport | 12.60 |
| A0A3B6LVY8 | CMP/dCMP-type deaminase domain-containing protein | Metal-binding | 14.90 |
| A0A3B6MXP5 | Cns1/TTC4 wheel domain-containing protein | - | 14.68 |
| W5G289 | Complex 1 LYR protein domain-containing protein | - | 12.63 |
| A0A3B6RMD0 | CRAL-TRIO domain-containing protein | - | 13.72 |
| A0A0R6UPC9 | CslF6 | Cell wall biogenesis/degradation | 14.93 |
| A0A3B6IQY2 | Cupin type-1 domain-containing protein | - | 16.53 |
| A0A3B6SF34 | Cyclic phosphodiesterase | - | 15.16 |
| A0A3B6HXU4 | Cytochrome P450 | Heme | 11.92 |
| A0A3B6SUC6 | Cytochrome P450 | Heme | 14.14 |
| A0A3B6RQ17 | Cytochrome P450 | Heme | 14.04 |
| W4ZVB1 | Defensin | Disulfide bond | 16.99 |
| A0A3B6JGH7 | Dehydrin | - | 14.99 |
| A0A3B6KB32 | Diacylglycerol kinase (DAG kinase) (EC 2.7.1.107) | ATP-binding | 13.68 |
| A0A3B6KCM1 | Dihydrolipoamide acetyltransferase component of pyruvate dehydrogenase complex (EC 2.3.1.-) | Acyltransferase | 15.20 |
| A0A3B6RQ45 | DNA polymerase alpha subunit B | DNA replication | 15.53 |
| A0A3B6PI01 | DNA polymerase V | Nucleus | 14.40 |
| A0A3B6ITA2 | DNA/RNA-binding protein Alba-like domain-containing protein | Nucleus | 14.22 |
| A0A3B6NMU8 | DRBM domain-containing protein | - | 11.59 |
| A0A3B6SCK5 | DUF7725 domain-containing protein | - | 12.74 |
| A0A3B6JI82 | E1 ubiquitin-activating enzyme (EC 6.2.1.45) | ATP-binding | 12.24 |
| A0A3B6I0B8 | Early nodulin | Membrane | 14.96 |
| A0A3B6PRD8 | EF-hand domain-containing protein | - | 13.52 |
| G8A4X7 | Ent-kaurenoic acid oxidase | Heme | 14.14 |
| A0A3B6KQP7 | Eukaryotic translation initiation factor 3 subunit C (eIF3c) | Coiled coil | 13.37 |
| A0A3B6HXG5 | Exocyst complex component Sec3 PIP2-binding N-terminal domain-containing protein | Coiled coil | 12.50 |
| A0A3B6NPQ7 | Exostosin GT47 domain-containing protein | Glycosyltransferase | 14.28 |
| A0A3B6ET22 | Exostosin GT47 domain-containing protein | Glycosyltransferase | 15.89 |
| A0A3B6SM28 | Expansin | Cell wall | 14.37 |
| A0A341UE42 | F-actin-capping protein subunit beta | Actin capping | 15.25 |
| A0A3B6KI63 | FAD dependent oxidoreductase domain-containing protein | FAD | 14.59 |
| A0A3B6IX04 | FAD-binding domain-containing protein | Monooxygenase | 13.07 |
| A0A3B6KCG3 | Fatty acid hydroxylase domain-containing protein | Endoplasmic reticulum | 13.86 |
| A0A3B6HQY8 | Fungal lipase-type domain-containing protein | - | 13.57 |
| B6UKM9 | Gamma gliadin-A3 (Gamma gliadin-A4) (Gamma-gliadin) | - | 18.13 |
| M9TE79 | Gamma gliadin-B1 (Gamma-gliadin 3) | - | 17.22 |
| A0A3B6N3H5 | GCF C-terminal domain-containing protein | - | 13.44 |
| A0A3B6SD22 | GDSL esterase/lipase | Lipid degradation | 16.22 |
| A0A3B5XY05 | GDSL esterase/lipase | Lipid degradation | 15.44 |
| A0A3B6KR49 | GH10 domain-containing protein | Carbohydrate metabolism | 15.56 |
| A0A3B6LTP4 | Glycosyltransferase (EC 2.4.1.-) | Glycosyltransferase | 14.83 |
| T1QDI5 | Glycosyltransferase (EC 2.4.1.-) | Glycosyltransferase | 15.50 |
| A0A3B6QHI6 | Glycosyltransferase 6 | Glycosyltransferase | 15.74 |
| A0A3B6QI93 | Glycosyltransferase 61 catalytic domain-containing protein | Glycoprotein | 15.34 |
| A0A3B6IWS5 | Gnk2-homologous domain-containing protein | Cell junction | 14.73 |
| A0A3B6NT82 | GPI-anchored protein | Membrane | 16.66 |
| A0A3B6PGB9 | GST C-terminal domain-containing protein | - | 15.97 |
| B2ZGG5 | GTPase SAR1 | Endoplasmic reticulum | 14.48 |
| Q9ZSR6 | Heat shock protein HSP26 (Small heat-shock protein) | - | 15.83 |
| A0A3B5XVV7 | Heparan-alpha-glucosaminide N-acetyltransferase catalytic domain-containing protein | Membrane | 14.93 |
| A0A3B6LU11 | Hexosyltransferase | Glycosyltransferase | 12.40 |
| A0A3B6NS03 | Hexosyltransferase (EC 2.4.1.-) | Glycosyltransferase | 14.40 |
| A0A3B6LWD9 | histidine kinase (EC 2.7.13.3) | Copper | 14.59 |
| A0A3B6IK88 | Histidine-containing phosphotransfer protein | Cytokinin signaling pathway | 17.03 |
| A0A3B6JEB3 | HMA domain-containing protein | Lipoprotein | 15.32 |
| A0A3B6RD87 | Homeobox domain-containing protein | DNA-binding | 12.74 |
| A0A3B6MV56 | inositol-1,3,4-trisphosphate 5/6-kinase (EC 2.7.1.159) | ATP-binding | 13.67 |
| W5CTA6 | Knottin scorpion toxin-like domain-containing protein | Disulfide bond | 17.44 |
| A0A3B6NUF5 | Laccase (EC 1.10.3.2) (Benzenediol:oxygen oxidoreductase) | Apoplast | 15.26 |
| A0A3B6MKI4 | leucine--tRNA ligase (EC 6.1.1.4) (Leucyl-tRNA synthetase) | Aminoacyl-tRNA synthetase | 12.60 |
| A0A3B6MPT7 | Metallo-beta-lactamase domain-containing protein | - | 14.83 |
| A0A3B5Z097 | Methyltransferase (EC 2.1.1.-) | Glycoprotein | 13.89 |
| A0A3B6NTW9 | Methyltransferase type 11 domain-containing protein | Methyltransferase | 12.14 |
| W5E953 | Mitochondrial import inner membrane translocase subunit | Chaperone | 15.53 |
| A0A3B6PQ92 | Mitochondrial import inner membrane translocase subunit | Chaperone | 13.44 |
| A0A3B5Z086 | MLO-like protein | Calmodulin-binding | 11.33 |
| A0A3B6NSM3 | monodehydroascorbate reductase (NADH) (EC 1.6.5.4) | FAD | 14.69 |
| A0A3B6NSV4 | MSP domain-containing protein | Coiled coil | 15.04 |
| A0A3B6KPW6 | Multiple inositol polyphosphate phosphatase 1 (EC 3.1.3.62) | Cell membrane | 14.76 |
| A0A3B5Y512 | Myb-like domain-containing protein | Chromatin regulator | 15.72 |
| A0A3B6PNQ5 | Myb-like domain-containing protein | - | 14.47 |
| A0A3B6RK39 | NAD-dependent epimerase/dehydratase domain-containing protein | Oxidoreductase | 14.82 |
| A0A3B6QM40 | Neprosin domain-containing protein | - | 13.12 |
| A0A3B6KG61 | OCRE domain-containing protein | - | 14.00 |
| A0A3B6MSP0 | O-fucosyltransferase family protein | Carbohydrate metabolism | 15.33 |
| A0A3B6LWF6 | Patatin (EC 3.1.1.-) | Hydrolase | 13.61 |
| A0A3B6QNR5 | PB1 domain-containing protein | - | 14.30 |
| A0A3B6IWU9 | PCI domain-containing protein | Proteasome | 14.29 |
| A0A3B6NR90 | PDZ domain-containing protein | - | 14.46 |
| A0A3B6T8A9 | Peptidase A1 domain-containing protein | Aspartyl protease | 16.78 |
| A0A3B6TQF4 | Peptidase A1 domain-containing protein | Aspartyl protease | 14.13 |
| A0A3B6SMA2 | Peptidase A1 domain-containing protein | Aspartyl protease | 14.58 |
| A0A3B6TR73 | Peptidase A1 domain-containing protein | Aspartyl protease | 16.25 |
| A0A3B6RQL6 | Peptidyl-prolyl cis-trans isomerase (PPIase) (EC 5.2.1.8) | Isomerase | 15.05 |
| A0A3B6HWT4 | PH domain-containing protein | Lipid transport | 14.06 |
| A0A3B6I1C1 | PHD finger protein ING | Chromatin regulator | 13.77 |
| A0A3B6KFM2 | PHD-type domain-containing protein | Metal-binding | 12.09 |
| A0A3B6JNZ1 | Phytocyanin domain-containing protein | Copper | 15.28 |
| A0A3B6MKQ2 | Plasma membrane ATPase (EC 7.1.2.1) | ATP-binding | 16.01 |
| A0A3B6JCH7 | Pre-rRNA-processing protein TSR2 homolog | - | 14.99 |
| A0A3B5XVP6 | Pre-rRNA-processing protein TSR2 homolog | - | 12.61 |
| J7HWD3 | Prolamin | - | 19.33 |
| K7XRA1 | Prolamin | - | 18.21 |
| A0A3B6NLR8 | Prolyl endopeptidase (EC 3.4.21.-) | Hydrolase | 13.92 |
| A0A3B6KKQ3 | Proteasome assembly chaperone 4 | - | 14.86 |
| A0A3B6SE72 | Protein kinase domain-containing protein | ATP-binding | 13.51 |
| A0A3B6RRX8 | Protein kinase domain-containing protein | ATP-binding | 12.34 |
| A0A3B6N0F6 | Protein NO VEIN C-terminal domain-containing protein | - | 13.04 |
| A0A3B6PSK7 | protein-serine/threonine phosphatase (EC 3.1.3.16) | Hydrolase | 16.30 |
| A0A3B6RJZ1 | Putative tRNA (cytidine(32)/guanosine(34)-2'-O)-methyltransferase (EC 2.1.1.205) | Cytoplasm | 14.41 |
| A0A3B6RRZ6 | PX domain-containing protein | - | 14.68 |
| A0A3B5XVE0 | RING-type domain-containing protein | Coiled coil | 13.98 |
| A0A3B6HSE4 | RING-type E3 ubiquitin transferase (EC 2.3.2.27) | Chaperone | 12.15 |
| A0A3B6NWX2 | RING-type E3 ubiquitin transferase (EC 2.3.2.27) | Metal-binding | 10.80 |
| A0A3B5XT84 | RING-type E3 ubiquitin transferase (EC 2.3.2.27) | DNA-binding | 14.81 |
| A0A3B6TTH2 | RNase III domain-containing protein | - | 14.58 |
| A0A3B6RFB3 | RRM domain-containing protein | - | 13.50 |
| A0A3B5XSQ8 | RRM domain-containing protein | - | 15.84 |
| A0A3B6QMR3 | Salt-induced YSK2 dehydrin 3 | - | 14.20 |
| A0A3B5Y552 | Saposin B-type domain-containing protein | Disulfide bond | 13.88 |
| A0A3B6QJ23 | SHSP domain-containing protein | - | 15.77 |
| A0A3B6HQJ3 | Small-subunit processome Utp12 domain-containing protein | Nucleus | 13.82 |
| A0A3B6NXG4 | SMP-LTD domain-containing protein | Endoplasmic reticulum | 14.00 |
| A0A3B6IWH2 | STAS domain-containing protein | Membrane | 13.59 |
| A0A3B6SC19 | Subtilisin-like protease | Glycoprotein | 14.16 |
| A0A3B6LU07 | Succinate dehydrogenase [ubiquinone] flavoprotein subunit, mitochondrial (EC 1.3.5.1) | Electron transport | 15.17 |
| A0A3B6GST0 | Survival protein SurE-like phosphatase/nucleotidase domain-containing protein | Hydrolase | 15.90 |
| A0A3B6I5D4 | Thromboxane-A synthase | Heme | 14.17 |
| A0A3B6ILK3 | Transcription factor CBF/NF-Y/archaeal histone domain-containing protein | - | 11.87 |
| A0A3B5Y593 | Transcription factor VOZ1 | - | 13.36 |
| A0A3B6KUR6 | Tubulin beta chain | Cytoplasm | 16.77 |
| A0A3B6LJG0 | U3 small nucleolar ribonucleoprotein protein MPP10 | Nucleus | 15.86 |
| A0A3B6TGT0 | Ufm1-specific protease | Hydrolase | 15.13 |
| A0A341PTZ7 | Uncharacterized protein | Membrane | 13.21 |
| A0A3B5XYE2 | Uncharacterized protein | - | 13.94 |
| A0A3B5YUF3 | Uncharacterized protein | Glycosyltransferase | 15.59 |
| A0A3B6I3X8 | Uncharacterized protein | - | 13.83 |
| A0A3B6PJF1 | Uncharacterized protein | Coiled coil | 11.77 |
| A0A3B6QKG1 | Uncharacterized protein | Membrane | 11.12 |
| A0A3B6RMN4 | Uncharacterized protein | - | 14.78 |
| A0A3B6SJX4 | Uncharacterized protein | Membrane | 16.31 |
| A0A1D5YAY8 | Uncharacterized protein | Membrane | 14.57 |
| A0A3B6KKC0 | Uncharacterized protein | - | 14.48 |
| A0A3B6NM71 | Uncharacterized protein | - | 14.27 |
| A0A3B6PJ71 | Uncharacterized protein | Membrane | 15.87 |
| A0A3B6QA75 | Uncharacterized protein | - | 13.99 |
| A0A3B6RMU7 | Uncharacterized protein | Membrane | 12.79 |
| A0A3B6TMH9 | Uncharacterized protein | Abscisic acid signaling pathway | 15.08 |
| A0A3B5Y051 | Uncharacterized protein | Membrane | 14.24 |
| A0A3B6I0W4 | Uncharacterized protein | Membrane | 16.07 |
| A0A3B6JM39 | Uncharacterized protein | ATP-binding | 15.30 |
| A0A3B6RGZ6 | Uncharacterized protein | Hydrolase | 14.53 |
| A0A3B6KBH6 | Uncharacterized protein | - | 14.87 |
| A0A3B6RJG8 | Uncharacterized protein | - | 14.60 |
| A0A3B6SFT3 | Uncharacterized protein | Membrane | 12.61 |
| A0A3B6NQA3 | Uncharacterized protein | - | 16.50 |
| A0A3B6RQD9 | Uncharacterized protein | DNA-binding | 14.11 |
| A0A3B6HZB7 | Uncharacterized protein | - | 16.34 |
| A0A3B5Y650 | Uncharacterized protein | Plant defense | 19.93 |
| A0A3B6RB66 | WAT1-related protein | Membrane | 13.78 |
| A0A3B6MYI7 | X8 domain-containing protein | - | 15.87 |
| A0A3B5Y4J9 | X8 domain-containing protein | - | 14.94 |
| A0A3B6IR79 | YTH domain-containing family protein | - | 12.93 |
| A0A3B6QEG6 | ZZ-type domain-containing protein | Cytoplasmic vesicle | 13.67 |

*Protein ID from UniProt

^#^Keyword and function derived from UniProt terms.

**Table S3: Prioritized core wheat proteins with defined roles in pathogen defense and mycotoxin detoxification.**

| **Protein Name** | **# proteins** | **Function** |
| --- | --- | --- |
| 14-3-3 domain-containing protein | 2 | protein localization; signal transduction |
| 3-methyl-2-oxobutanoate dehydrogenase (2-methylpropanoyl-transferring) | 1 | branched-chain amino acid catabolic process; response to nutrient |
| 5-oxoprolinase | 1 | glutathione metabolic process |
| Aldehyde oxidase | 1 |  |
| Cytochrome P450 | 23 | lipid metabolic process |
| Globin domain-containing protein | 1 |  |
| Glutathione reductase (GRase) | 4 | cell redox homeostasis; glutathione metabolic process |
| Glutathione S-transferase | 26 | glutathione metabolic process |
| Glutenin, high molecular weight subunit DX5 | 1 | carbohydrate storage |
| glycerol kinase (ATP:glycerol 3-phosphotransferase) | 1 | glycerol catabolic process; glycerol metabolic process; glycerol-3-phosphate biosynthetic process; triglyceride metabolic process |
| Glycosyltransferase | 13 |  |
| Histone-binding protein RBBP4 N-terminal domain-containing protein | 3 | chromatin remodeling; regulation of DNA-templated transcription |
| Microsomal glutathione S-transferase 3 | 1 | leukotriene metabolic process |
| Monocopper oxidase-like protein SKU5 | 1 |  |
| Obtusifoliol 14-alpha demethylase | 1 | sterol biosynthetic process |
| P450 | 1 |  |
| Pathogenesis-related protein 1-1 | 1 | sexual reproduction |
| Phosphoinositide phospholipase C | 1 | intracellular signal transduction; lipid catabolic process |
| Protein arginine N-methyltransferase | 1 | methylation; regulation of DNA-templated transcription |
| Respiratory burst oxidase homolog I | 1 |  |
| Thromboxane-A synthase | 1 |  |
| Tricetin 3',4',5'-O-trimethyltransferase (TaOMT2) (Caffeic acid 3-O-methyltransferase) (TaCM) (Flavone O-methyltransferase 2) | 1 | biosynthetic process; flavonoid biosynthetic process; lignin biosynthetic process; methylation |
| Uncharacterized protein | 1 | COPII-coated vesicle budding; mRNA transport; protein import into nucleus |
| Xanthine dehydrogenase | 1 |  |
| Chitinase | 1 | cell wall macromolecule catabolic process; chitin catabolic process; defense response to fungus; polysaccharide catabolic process |
| Cinnamyl alcohol dehydrogenase | 1 | lignin biosynthetic process |
| Cinnamyl alcohol dehydrogenase | 1 | lignin biosynthetic process |
| Cinnamyl alcohol dehydrogenase | 1 | lignin biosynthetic process |
| Cinnamyl alcohol dehydrogenase | 1 | lignin biosynthetic process |
| Leucine-rich repeat-containing N-terminal plant-type domain-containing protein | 1 |  |
| Thaumatin-like protein | 1 | defense response |
| Thaumatin-like protein | 1 | defense response |

*Protein ID from UniProt

^#^Difference based on normalized LFQ intensities of proteins upon designated chemotype treatment. Log_2_ transformed data.

**Table S4: Fungal proteins exclusively detected upon 15ADON/3ANX chemotype treatment**

| **Protein ID*** | **Protein name** | **Keyword/**  **Function^#^** | **LFQ Intensity (log2)** |
| --- | --- | --- | --- |
| A0A098DP61 | Chromosome 2 | Membrane | 16.63 |
| A0A0E0RWF0 | Chromosome 1 | FAD | 15.72 |
| A0A0E0S0J1 | Chromosome 2 | Metal-binding | 14.69 |
| A0A0E0S9A4 | Chromosome 4 | - | 13.68 |
| A0A1C3YI55 | Chromosome 1 | LIM domain | 13.90 |
| A0A1C3YKE6 | Large ribosomal subunit protein bL32m | Mitochondrion | 13.39 |
| A0A1C3YL34 | DNA topoisomerase I | DNA-binding | 14.20 |
| A0A1C3YM78 | Chromosome 4 | Nucleus | 16.27 |
| I1R9X4 | Chromosome 1 | - | 14.80 |
| I1RC17 | Chromosome 1 | - | 15.00 |
| I1RER3 | Chromosome 1 | - | 15.52 |
| I1RHN1 | Chromosome 2 | Metal-binding | 13.60 |
| I1RIL7 | Chromosome 2 | - | 14.65 |
| I1RIM6 | Chromosome 2 | - | 14.20 |
| I1RJP7 | Chromosome 2 | Hydrolase | 15.91 |
| I1RKI8 | Chromosome 2 | Membrane | 14.23 |
| I1RKN7 | Chromosome 2 | NADP | 14.79 |
| I1RNQ3 | Chromosome 3 | - | 15.94 |
| I1RPC6 | Trafficking protein particle complex subunit | Endoplasmic reticulum | 12.34 |
| I1RPX0 | Chromosome 3 | - | 13.60 |
| I1RTD5 | Chromosome 4 | - | 15.09 |
| I1RY34 | Chromosome 4 | Oxidoreductase | 13.57 |
| I1RZ36 | Sensitive to high expression protein 9, mitochondrial | Coiled coil | 14.03 |
| I1RZA5 | ER membrane protein complex subunit 4 | Endoplasmic reticulum | 14.95 |
| I1S060 | U3 small nucleolar RNA-associated protein 25 | Nucleus | 13.78 |
| I1S2C2 | Chromosome 3 | - | 14.73 |
| I1S3S8 | Chromosome 3 | - | 14.45 |
| I1SA47 | Chromosome 1 | - | 14.67 |
| Q4I4A4 | Rhomboid-type serine protease 2 (EC 3.4.21.105) | Golgi apparatus | 15.51 |
| Q4I6B0 | Mitochondrial import inner membrane translocase subunit TIM13 | Chaperone | 14.14 |
| Q4IJW4 | Mitochondrial import inner membrane translocase subunit TIM8 | Chaperone | 15.39 |
| V6QV38 | Transcription initiation factor TFIID subunit 9 | Nucleus | 14.08 |
| A0A098E587 | Chromosome 3 | Membrane | 15.35 |
| A0A0E0RUQ7 | Chromosome 1 | mRNA processing | 15.24 |
| A0A0E0S0X2 | Chromosome 2 | - | 14.72 |
| A0A1C3YIL9 | Mannosyltransferase | Endoplasmic reticulum | 13.30 |
| I1RBU8 | Chromosome 1 | - | 15.34 |
| I1RC24 | Chromosome 1 | Nucleus | 15.29 |
| I1RC94 | Chromosome 1 | - | 14.39 |
| I1RD00 | Chromosome 1 | - | 14.14 |
| I1RDZ4 | Ubiquitin carboxyl-terminal hydrolase | Hydrolase | 14.35 |
| I1RE02 | Chromosome 1 | - | 15.43 |
| I1REV2 | Chromosome 1 | Hydrolase | 15.13 |
| I1RG58 | Chromosome 1 | Purine metabolism | 14.55 |
| I1RHW3 | ribonuclease T2 | Disulfide bond | 14.99 |
| I1RJ46 | Chromosome 2 | Membrane | 14.00 |
| I1RJF8 | Chromosome 2 | Aspartyl protease | 16.65 |
| I1RJG8 | Chromosome 2 | - | 15.91 |
| I1RN23 | Chromosome 3 | Membrane | 13.82 |
| I1RNP1 | Chromosome 3 | GTP-binding | 14.90 |
| I1RQC3 | Chromosome 3 | Membrane | 13.80 |
| I1RR61 | Chromosome 4 | Nucleus | 14.19 |
| I1RSS1 | Derlin | Endoplasmic reticulum | 16.21 |
| I1RTD7 | Chromosome 4 | - | 15.16 |
| I1RV22 | Putative acetyltransferase FG08082 | Acyltransferase | 15.16 |
| I1RWN4 | Chromosome 2 | GTP-binding | 14.36 |
| I1RXR5 | Chromosome 4 | - | 14.57 |
| I1RYE5 | Chromosome 4 | Membrane | 14.71 |
| I1RYQ3 | NADH dehydrogenase [ubiquinone] 1 beta subcomplex subunit 11, mitochondrial | Electron transport | 15.18 |
| I1RZA4 | Chromosome 4 | Cell membrane | 12.67 |
| I1RZK8 | Chromosome 4 | Pyridoxal phosphate | 15.08 |
| I1RZP0 | 5-hydroxyisourate hydrolase | Hydrolase | 14.74 |
| I1S073 | Chromosome 1 | - | 14.20 |
| I1S1J0 | Chromosome 1 | Hydrolase | 15.44 |
| I1S2P6 | Chromosome 3 | - | 14.98 |
| I1S522 | Chromosome 1 | Membrane | 14.05 |
| Q4I6A4 | Histone acetyltransferase type B catalytic subunit | Acyltransferase | 15.39 |
| Q4IP34 | Pre-mRNA-processing ATP-dependent RNA helicase PRP5 | ATP-binding | 13.52 |
| V6R5Q6 | alpha-1,2-Mannosidase | Calcium | 14.34 |
| A0A098DJZ7 | Chromosome 2 | Membrane | 14.15 |
| A0A098E3X0 | Chromosome 3 | ANK repeat | 14.61 |
| A0A0E0RN05 | Chromosome 1 | Nucleus | 15.84 |
| A0A0E0S067 | Chromosome 2 | - | 16.51 |
| A0A0E0SA11 | Chromosome 4 | - | 16.06 |
| A0A0E0SDV0 | non-specific serine/threonine protein kinase | ATP-binding | 13.89 |
| A0A0E0SKJ2 | ER membrane protein complex subunit 3 | Membrane | 14.96 |
| A0A1C3YJ33 | Carboxypeptidase | Apoptosis | 15.03 |
| A0A1C3YKU0 | Chromosome 4 | - | 14.70 |
| A0A1C3YL89 | Chromosome 4 | GTP-binding | 15.68 |
| A0A1C3YLI5 | Chromosome 4 | ATP-binding | 14.02 |
| I1RA11 | Chromosome 1 | - | 14.85 |
| I1RA63 | Chromosome 1 | Exonuclease | 14.61 |
| I1RBC7 | DNA-directed RNA polymerase subunit beta | DNA-directed RNA polymerase | 13.76 |
| I1RC20 | Translation machinery-associated protein 22 | Cytoplasm | 14.56 |
| I1RC82 | Chromosome 1 | Coiled coil | 13.44 |
| I1RDJ4 | Chromosome 1 | - | 14.12 |
| I1RFC0 | Chromosome 1 | - | 13.70 |
| I1RH38 | Chromosome 2 | - | 15.06 |
| I1RHB9 | Chromosome 2 | - | 16.86 |
| I1RKU0 | Chromosome 2 | Copper | 14.81 |
| I1RM25 | RNA helicase | ATP-binding | 13.42 |
| I1RMH4 | Ceramide very long chain fatty acid hydroxylase | Endoplasmic reticulum | 15.24 |
| I1RMW4 | Ferrochelatase | Heme biosynthesis | 15.54 |
| I1RN50 | Chromosome 3 | Membrane | 14.65 |
| I1RNL7 | Signal peptidase complex subunit 2 | Endoplasmic reticulum | 15.91 |
| I1RNM4 | Alpha-1,3-glucosyltransferase | Endoplasmic reticulum | 13.18 |
| I1RPA0 | Chromosome 3 | Metal-binding | 14.88 |
| I1RQ85 | Large ribosomal subunit protein uL23m | - | 12.82 |
| I1RQJ9 | Chromosome 3 | - | 13.96 |
| I1RRC5 | Chromosome 4 | - | 12.83 |
| I1RRU2 | non-specific serine/threonine protein kinase | ATP-binding | 13.82 |
| I1RRV9 | Chromosome 4 | Membrane | 15.68 |
| I1RS87 | Chromosome 4 | Hydrolase | 15.81 |
| I1RSH3 | Chromosome 4 | Golgi apparatus | 14.71 |
| I1RSQ1 | Chromosome 4 | - | 13.26 |
| I1RTS2 | Chromosome 4 | - | 14.82 |
| I1RTZ8 | Carboxypeptidase M14B (Carboxypeptidase MCPB) | - | 14.22 |
| I1RVM1 | Chromosome 2 | - | 14.45 |
| I1RWJ6 | Ubiquitin carboxyl-terminal hydrolase | Hydrolase | 15.17 |
| I1RWU9 | Protein YIF1 | Endoplasmic reticulum | 12.75 |
| I1RXF5 | Synaptobrevin homolog YKT6 | Coiled coil | 13.99 |
| I1RZ99 | Chromosome 4 | - | 14.71 |
| I1RZB5 | DNA-directed RNA polymerases I, II, and III subunit RPABC1 | Nucleus | 14.81 |
| I1S0N1 | Chromosome 1 | Membrane | 15.10 |
| I1S1U2 | Chromosome 1 | Hydrolase | 15.95 |
| I1S238 | Prefoldin subunit 4 | Chaperone | 15.60 |
| I1S7S4 | Chromosome 3 | - | 13.62 |
| I1S8F0 | Chromosome 4 | Membrane | 15.17 |
| I1S9Z3 | Chromosome 1 | Membrane | 15.38 |
| Q4I375 | Mitochondrial import inner membrane translocase subunit TIM16 | Membrane | 14.69 |
| Q4IPH4 | Peptidyl-prolyl cis-trans isomerase H | Isomerase | 15.95 |
| V6RH79 | Chromosome 4 | - | 15.91 |
| V6RM02 | Glycine cleavage system H protein | Lipoyl | 15.30 |
| A0A098D6N4 | Cutinase | Disulfide bond | 13.98 |
| A0A098DIB1 | Chromosome 2 | - | 15.02 |
| A0A098DMM9 | Splicing factor U2AF subunit | mRNA processing | 15.41 |
| A0A098DXD6 | Pyruvate decarboxylase | Decarboxylase | 15.03 |
| A0A098E017 | Chromosome 3 | Membrane | 14.15 |
| A0A0E0SDN6 | Chromosome 4 | - | 14.40 |
| A0A1C3YI81 | Chromosome 1 | mRNA processing | 13.48 |
| A0A1C3YIQ7 | Chromosome 1 | Coiled coil | 14.16 |
| A0A1C3YIZ4 | Chromosome 1 | - | 14.29 |
| A0A1C3YKY8 | Chromosome 4 | DNA-binding | 13.77 |
| A0A1C3YLU6 | Chromosome 4 | CBS domain | 14.99 |
| A0A1C3YMX9 | Chromosome 2 | Iron | 15.47 |
| A0A1C3YMY3 | Chromosome 2 | - | 15.66 |
| I1R9C0 | Chromosome 1 | - | 14.81 |
| I1R9X7 | Chromosome 1 | Nucleus | 13.65 |
| I1RA50 | Chromosome 1 | Membrane | 13.80 |
| I1RAK7 | Chromosome 1 | Endoplasmic reticulum | 15.63 |
| I1RB41 | Chromosome 1 | - | 16.09 |
| I1RBA3 | Chromosome 1 | - | 14.20 |
| I1RBD7 | Chromosome 1 | - | 15.07 |
| I1RCQ3 | Chromosome 1 | Chaperone | 15.59 |
| I1RD53 | Chromosome 1 | - | 14.88 |
| I1RFV4 | Chromosome 1 | - | 14.25 |
| I1RG29 | Chromosome 1 | Pyridoxal phosphate | 13.72 |
| I1RH82 | Chromosome 2 | Carbohydrate metabolism | 15.08 |
| I1RHL4 | Chromosome 2 | - | 15.40 |
| I1RIF2 | Chromosome 2 | - | 15.44 |
| I1RJH8 | Signal peptidase complex catalytic subunit SEC11 | Endoplasmic reticulum | 14.87 |
| I1RKE7 | Chromosome 2 | - | 15.35 |
| I1RLB7 | Chromosome 3 | Copper | 15.83 |
| I1RLM3 | Kynureninase | Cytoplasm | 15.42 |
| I1RMF6 | Chromosome 3 | - | 14.37 |
| I1RMV5 | U2 small nuclear ribonucleoprotein A' | Leucine-rich repeat | 14.40 |
| I1RN81 | Chromosome 3 | Cell membrane | 15.37 |
| I1RPN8 | Chromosome 3 | ATP-binding | 14.16 |
| I1RQF2 | Prefoldin subunit 3 | Chaperone | 15.53 |
| I1RRE2 | Pre-rRNA-processing protein RIX1 | Nucleus | 13.49 |
| I1RRW3 | Chromosome 4 | Metal-binding | 16.21 |
| I1RSV9 | Chromosome 4 | - | 15.01 |
| I1RXT3 | Chromosome 4 | Membrane | 14.85 |
| I1RYA2 | D-malate dehydrogenase | Magnesium | 14.06 |
| I1RYP7 | Chromosome 4 | Golgi apparatus | 14.72 |
| I1S053 | Chromosome 1 | Nucleus | 15.91 |
| I1S1V1 | Chromosome 3 | - | 15.49 |
| I1S227 | Chromosome 3 | Membrane | 14.63 |
| I1S2J3 | Short chain dehydrogenase FGM9 | NADP | 13.88 |
| I1S2X2 | Chromosome 3 | Membrane | 15.93 |
| I1S7A8 | Chromosome 3 | - | 14.89 |
| I1S9T8 | Chromosome 4 | - | 15.09 |
| I1SAR6 | Chromosome 3 | - | 15.21 |
| Q4I1T9 | Protein EFR3 | - | 17.59 |
| Q4IPZ1 | Mitochondrial import inner membrane translocase subunit TIM10 | Chaperone | 15.00 |
| V6QXN0 | Chromosome 1 | - | 14.58 |
| A0A098DKM1 | Chromosome 2 | GTP-binding | 12.57 |
| A0A098DPS3 | Chromosome 4 | GTP-binding | 15.36 |
| A0A098DUL9 | Chromosome 4 | - | 15.08 |
| A0A0E0RU86 | Chromosome 1 | - | 13.21 |
| A0A0E0SH52 | dTMP kinase | Kinase | 14.97 |
| A0A1C3YHC8 | Chromosome 1 | - | 15.07 |
| A0A1C3YIL2 | Chromosome 1 | Membrane | 15.01 |
| A0A1C3YIN9 | Chromosome 1 | Chaperone | 13.67 |
| A0A1C3YL14 | Chromosome 4 | NADP | 14.01 |
| A0A1C3YLK8 | Chromosome 4 | - | 15.65 |
| A0A1C3YMA0 | Chromosome 2 | - | 14.44 |
| A0A1C3YMD5 | Chromosome 2 | Lipid degradation | 14.73 |
| A0A1C3YMY7 | Chromosome 2 | ATP-binding | 14.73 |
| A0A1C3YN57 | Chromosome 2 | Nucleotide-binding | 15.11 |
| I1RAB3 | Chromosome 1 | Magnesium | 14.88 |
| I1RC67 | Chromosome 1 | - | 13.68 |
| I1RC81 | Chromosome 1 | Exonuclease | 14.67 |
| I1RDC7 | Cellulase | Glycosidase | 15.09 |
| I1RDD4 | Chromosome 1 | - | 15.26 |
| I1RDU6 | Chromosome 1 | Lyase | 15.53 |
| I1RDV7 | Chromosome 1 | - | 15.40 |
| I1REK7 | Chromosome 1 | - | 15.47 |
| I1RHJ2 | Glycerol-3-phosphate dehydrogenase | FAD | 13.94 |
| I1RHZ7 | Chromosome 2 | FAD | 15.34 |
| I1RIX2 | Carboxypeptidase | Carboxypeptidase | 15.20 |
| I1RJP4 | Translation machinery-associated protein 20 | Cytoplasm | 16.07 |
| I1RK03 | Pantoate--beta-alanine ligase | ATP-binding | 15.32 |
| I1RKC6 | Chromosome 2 | Nucleus | 14.66 |
| I1RLV1 | Chromosome 3 | Membrane | 14.88 |
| I1RMB2 | Chromosome 3 | Membrane | 14.43 |
| I1RMG4 | Chromosome 3 | Lipid metabolism | 13.59 |
| I1RMK6 | Selenoprotein O | ATP-binding | 14.15 |
| I1RNR3 | Chromosome 3 | - | 14.68 |
| I1RQM6 | H/ACA ribonucleoprotein complex subunit | Nucleus | 16.03 |
| I1RRB8 | Chromosome 4 | Amino-acid transport | 13.07 |
| I1RWJ5 | Chromosome 2 | - | 12.00 |
| I1RX50 | Autophagy-related protein 3 | Autophagy | 14.28 |
| I1RX63 | Chromosome 2 | - | 15.96 |
| I1RXS0 | Chromosome 4 | - | 13.63 |
| I1RY41 | Chromosome 4 | - | 13.91 |
| I1RYZ4 | Chromosome 4 | Membrane | 16.48 |
| I1RZ94 | Chromosome 4 | DNA replication | 15.43 |
| I1RZJ5 | Signal recognition particle subunit SRP14 (Signal recognition particle 14 kDa protein) | Cytoplasm | 15.62 |
| I1S027 | Chromosome 1 | Membrane | 15.00 |
| I1S0J7 | Ubiquitin-like modifier-activating enzyme ATG7 | Autophagy | 14.54 |
| I1S0V8 | Chromosome 1 | - | 13.34 |
| I1S250 | Chromosome 3 | - | 14.30 |
| I1S357 | Chromosome 3 | - | 16.44 |
| Q4IJH1 | ATP-dependent RNA helicase DBP3 | ATP-binding | 14.83 |
| V6R5P0 | Chromosome 2 | Metal-binding | 15.10 |
| V6RVW7 | Chromosome 3 | Heme | 13.42 |
| A0A098DRL8 | Chromosome 4 | Glycoprotein | 14.83 |
| A0A098DTV2 | Chromosome 4 | - | 15.49 |
| A0A0E0SEM1 | Chromosome 4 | CBS domain | 14.68 |
| A0A0E0SPV0 | Chromosome 3 | ATP-binding | 14.54 |
| A0A1C3YIB7 | Chromosome 1 | - | 14.67 |
| A0A1C3YLA4 | Chromosome 4 | - | 15.15 |
| A0A1C3YMW0 | Chromosome 2 | - | 15.88 |
| A0A1C3YMW3 | Chromosome 2 | Iron | 14.56 |
| A0A1C3YN49 | Chromosome 2 | - | 14.95 |
| A0A1C3YNA2 | Chromosome 2 | - | 15.59 |
| I1RAG4 | Chromosome 1 | - | 15.82 |
| I1RB05 | Chromosome 1 | - | 14.57 |
| I1RCX0 | Chromosome 1 | Membrane | 15.12 |
| I1RD12 | Chromosome 1 | Membrane | 13.96 |
| I1RDK8 | Chromosome 1 | - | 14.21 |
| I1RJC4 | Chromosome 2 | NADP | 15.46 |
| I1RJS1 | Chromosome 2 | Ligase | 15.42 |
| I1RP46 | Chromosome 3 | - | 13.96 |
| I1RQE2 | Elicitor of plant defense protein 1 | Metal-binding | 13.34 |
| I1RTP1 | Chromosome 4 | Chromophore | 13.32 |
| I1RW00 | 3,4-dihydroxy-2-butanone 4-phosphate synthase (EC 4.1.99.12) | Lyase | 15.28 |
| I1RWD2 | aldehyde dehydrogenase (NAD(+)) (EC 1.2.1.3) | Oxidoreductase | 13.82 |
| I1RWE1 | Casein kinase II subunit beta (CK II beta) | - | 15.47 |
| I1RYY5 | COP9 signalosome complex subunit 6 | Cytoplasm | 14.91 |
| I1RZX9 | Chromosome 1 | GTP-binding | 14.04 |
| I1S069 | Chromosome 1 | - | 14.75 |
| I1S0M4 | Chromosome 1 | Iron | 16.27 |
| I1S1Q8 | Chromosome 1 | - | 14.83 |
| Q4I5I4 | Transcription elongation factor SPT5 | mRNA processing | 14.86 |
| A0A098D7T0 | Chromosome 1 | Membrane | 14.97 |
| A0A098DDV4 | 3-isopropylmalate dehydrogenase (EC 1.1.1.85) | Amino-acid biosynthesis | 14.97 |
| A0A098DQB4 | Chromosome 4 | ATP-binding | 14.87 |
| A0A098DS79 | Glutathione hydrolase (EC 2.3.2.2) | Acyltransferase | 15.65 |
| A0A098DTS4 | Chromosome 4 | - | 15.92 |
| A0A098E015 | Chromosome 3 | Acyltransferase | 16.37 |
| A0A0E0RTW2 | Amine oxidase (EC 1.4.3.-) | Copper | 14.09 |
| A0A0E0S6R7 | Chromosome 2 | - | 13.20 |
| A0A0E0S7B3 | Chromosome 2 | - | 14.45 |
| A0A0E0SIF3 | Chromosome 3 | - | 13.71 |
| A0A1C3YI31 | ubiquitinyl hydrolase 1 (EC 3.4.19.12) | Hydrolase | 14.89 |
| A0A1C3YJZ9 | Chromosome 3 | - | 15.46 |
| A0A1C3YK02 | trimethyllysine dioxygenase (EC 1.14.11.8) | Carnitine biosynthesis | 15.33 |
| A0A1C3YKW7 | Small ribosomal subunit protein mS29 | Mitochondrion | 14.59 |
| A0A1C3YL67 | Chromosome 4 | - | 15.64 |
| A0A1C3YL82 | Chromosome 4 | Amino-acid biosynthesis | 13.33 |
| I1RAD1 | U6 snRNA-associated Sm-like protein LSm6 | Cytoplasm | 15.19 |
| I1RB49 | Chromosome 1 | - | 16.12 |
| I1RBF8 | Chromosome 1 | - | 14.88 |
| I1RCB3 | Nuclear protein localization protein 4 | mRNA transport | 15.84 |
| I1RCY0 | Serine/threonine-protein phosphatase (EC 3.1.3.16) | Hydrolase | 15.16 |
| I1RDX6 | Chromosome 1 | - | 13.20 |
| I1REI0 | Chromosome 1 | Hydrolase | 15.34 |
| I1RFB7 | Pectate lyase (EC 4.2.2.2) | Calcium | 15.32 |
| I1RIG2 | Chromosome 2 | - | 14.58 |
| I1RIX3 | Chromosome 2 | - | 15.38 |
| I1RJL6 | Chromosome 2 | - | 14.62 |
| I1RJL8 | Chromosome 2 | - | 15.51 |
| I1RK12 | Chromosome 2 | Membrane | 16.72 |
| I1RM77 | Phosphoinositide phospholipase C (EC 3.1.4.11) | Hydrolase | 14.29 |
| I1RMV1 | Transcription initiation factor IIA subunit 2 | Nucleus | 14.99 |
| I1RPC0 | Chromosome 3 | Membrane | 14.36 |
| I1RSH0 | Chromosome 4 | Chaperone | 15.84 |
| I1RSR3 | Chromosome 4 | GTP-binding | 14.76 |
| I1RW32 | Chromosome 2 | Cytoplasm | 14.91 |
| I1RW89 | Chromosome 2 | ANK repeat | 15.95 |
| I1RX06 | Chromosome 2 | - | 15.40 |
| I1S074 | Vacuolar-sorting protein SNF7 | Endosome | 13.94 |
| I1S098 | Ubiquitin-conjugating enzyme E2 2 | ATP-binding | 13.97 |
| I1S223 | Chromosome 3 | FAD | 14.68 |
| I1S457 | Chromosome 2 | - | 16.28 |
| I1S4I7 | Chromosome 1 | - | 15.88 |
| I1SAM5 | Chromosome 3 | Coiled coil | 14.90 |
| I1SAX9 | Chromosome 2 | - | 16.03 |
| V6R0A3 | Chromosome 1 | NADP | 17.15 |
| V6RGB5 | Pyrroline-5-carboxylate reductase (EC 1.5.1.2) | Amino-acid biosynthesis | 15.73 |
| A0A098D991 | Chromosome 1 | Heme | 13.70 |
| A0A098DIG8 | Chromosome 2 | - | 15.35 |
| A0A098DLP1 | Chromosome 2 | - | 16.38 |
| A0A0E0S355 | Chromosome 2 | Membrane | 14.81 |
| A0A1C3YJI2 | Chromosome 3 | ATP-binding | 14.19 |
| A0A1C3YJK9 | Chromosome 1 | Cell cycle | 15.20 |
| A0A1C3YJR3 | DNA-directed RNA polymerase subunit beta (EC 2.7.7.6) | DNA-directed RNA polymerase | 14.30 |
| A0A1C3YMR3 | Chromosome 2 | - | 15.37 |
| I1RA04 | Chromosome 1 | - | 15.14 |
| I1RBM1 | Chromosome 1 | - | 16.68 |
| I1RD52 | Large ribosomal subunit protein uL3m | Coiled coil | 14.16 |
| I1RD82 | Autophagy-related protein 27 | Autophagy | 14.34 |
| I1RDE7 | chorismate synthase (EC 4.2.3.5) | Amino-acid biosynthesis | 14.33 |
| I1RGY6 | Chromosome 2 | - | 14.90 |
| I1RIV3 | Chromosome 2 | - | 15.93 |
| I1RJA9 | Chromosome 2 | Cell wall biogenesis/degradation | 14.56 |
| I1RJG2 | Inclusion body clearance protein IML2 | Membrane | 15.89 |
| I1RKL2 | Chromosome 2 | Bromodomain | 14.82 |
| I1RKX6 | Chromosome 2 | - | 15.73 |
| I1RLG9 | Chromosome 3 | - | 16.00 |
| I1RPE5 | Mevalonate kinase ERG12 (EC 2.7.1.36) | ATP-binding | 14.15 |
| I1RQ48 | Acyl-CoA desaturase (EC 1.14.19.1) | Electron transport | 15.42 |
| I1RQ88 | Chromosome 3 | Nucleus | 14.54 |
| I1RQA2 | Chromosome 3 | Methyltransferase | 14.64 |
| I1RRR5 | Signal peptidase subunit 3 | Endoplasmic reticulum | 15.67 |
| I1RSK5 | Chromosome 4 | FAD | 14.97 |
| I1RXT2 | Sorting nexin-4 (Autophagy-related protein 24) | Autophagy | 14.46 |
| I1RY32 | Peroxisomal membrane protein PEX14 (Peroxin-14) | Coiled coil | 15.74 |
| I1RYJ6 | Chromosome 4 | - | 15.30 |
| I1S1T8 | galacturonan 1,4-alpha-galacturonidase (EC 3.2.1.67) | Cell wall biogenesis/degradation | 14.68 |
| I1S3L9 | Chromosome 3 | - | 15.97 |
| A0A098DCZ3 | Beta-xylanase (EC 3.2.1.8) | Carbohydrate metabolism | 15.06 |
| A0A098DRI1 | Guanine nucleotide-exchange factor SEC12 | Endoplasmic reticulum | 15.10 |
| A0A098E4A1 | Chromosome 3 | - | 14.59 |
| A0A0E0S8X8 | Chromosome 4 | - | 15.04 |
| A0A1C3YI11 | Chromosome 1 | - | 14.62 |
| A0A1C3YIP4 | Sugar phosphate phosphatase (EC 3.1.3.-) | Hydrolase | 14.55 |
| A0A1C3YJ40 | Chromosome 1 | - | 15.29 |
| A0A1C3YMD0 | Chromosome 2 | Autocatalytic cleavage | 15.40 |
| I1RB50 | Small nuclear ribonucleoprotein G (snRNP-G) | mRNA processing | 16.07 |
| I1RBI5 | Chromosome 1 | - | 12.31 |
| I1RC68 | Chromosome 1 | - | 13.75 |
| I1RCN9 | Velvet complex subunit B | Cytoplasm | 15.99 |
| I1RE58 | cysteine-S-conjugate beta-lyase (EC 4.4.1.13) | Amino-acid biosynthesis | 15.17 |
| I1RH16 | Chromosome 2 | - | 14.30 |
| I1RK00 | Chromosome 2 | - | 14.76 |
| I1RLZ7 | Chromosome 3 | FAD | 14.60 |
| I1RQV1 | Chromosome 4 | Glycosidase | 15.09 |
| I1RRE0 | Exportin-T (Exportin(tRNA)) (tRNA exportin) | Cytoplasm | 15.61 |
| I1RRM3 | Large ribosomal subunit protein mL46 | Mitochondrion | 14.64 |
| I1RSY1 | Glutathione synthetase (GSH-S) (EC 6.3.2.3) | ATP-binding | 14.90 |
| I1RUE3 | Chromosome 4 | - | 15.08 |
| I1RUI4 | Chromosome 4 | - | 15.32 |
| I1RUM6 | Chromosome 4 | - | 15.11 |
| I1RUX8 | Chromosome 2 | - | 18.82 |
| I1RVX8 | Electron transfer flavoprotein-ubiquinone oxidoreductase (EC 1.5.5.1) | Electron transport | 14.88 |
| I1RZX3 | Chromosome 1 | Fatty acid metabolism | 16.00 |
| V6RHX2 | Chromosome 3 | Isomerase | 15.42 |
| A0A098D6R7 | Chromosome 1 | NAD | 15.94 |
| A0A098DDG2 | Chromosome 2 | - | 15.62 |
| A0A098DK18 | Chromosome 2 | Hydrolase | 15.31 |
| A0A098DLB3 | Chromosome 2 | - | 13.97 |
| A0A098DVM8 | Chromosome 3 | Golgi apparatus | 13.80 |
| A0A0E0S9Y6 | Chromosome 4 | Membrane | 15.41 |
| A0A1C3YK39 | Chromosome 3 | Metal-binding | 15.16 |
| I1RAB7 | Chromosome 1 | Calcium | 14.80 |
| I1RAW0 | Chromosome 1 | - | 14.38 |
| I1RBD6 | Serine/threonine-protein phosphatase (EC 3.1.3.16) | Cytoplasm | 14.12 |
| I1RBR4 | Sterol 14-alpha demethylase CYP51A (EC 1.14.14.154) | Endoplasmic reticulum | 15.81 |
| I1RCC5 | Chromosome 1 | Membrane | 14.96 |
| I1RCJ2 | ethanolamine kinase (EC 2.7.1.82) | - | 15.38 |
| I1RD38 | Chromosome 1 | - | 13.78 |
| I1RD63 | Vacuolar protein sorting-associated protein 29 | Protein transport | 14.59 |
| I1REY8 | Chromosome 1 | NADP | 15.53 |
| I1RFI9 | Chromosome 1 | Pyridoxal phosphate | 16.49 |
| I1RGF2 | Serine-threonine kinase receptor-associated protein | mRNA processing | 14.85 |
| I1RHL9 | Amine oxidase (EC 1.4.3.-) | Copper | 14.78 |
| I1RJY4 | Chromosome 2 | Cytoplasm | 13.55 |
| I1RLX1 | Chromosome 3 | - | 14.17 |
| I1RLZ5 | Chromosome 3 | - | 14.48 |
| I1RMH5 | Chromosome 3 | - | 14.69 |
| I1RMY3 | Chromosome 3 | Coiled coil | 13.66 |
| I1RN41 | Chromosome 3 | Membrane | 14.80 |
| I1RQK5 | Chromosome 3 | Golgi apparatus | 13.90 |
| I1RUE8 | Chromosome 4 | Oxidoreductase | 16.28 |
| I1RZJ9 | Chromosome 4 | Membrane | 14.51 |
| I1S1U7 | Chromosome 3 | Coiled coil | 14.15 |
| I1S3T1 | Chromosome 3 | - | 15.88 |
| I1S6P5 | Aspartate aminotransferase (EC 2.6.1.1) | Aminotransferase | 15.25 |
| A0A098D8U5 | Carboxylic ester hydrolase (EC 3.1.1.-) | Calcium | 13.90 |
| A0A098E5D2 | Chromosome 3 | - | 14.94 |
| A0A0E0S9X0 | Chromosome 4 | - | 15.27 |
| A0A0E0SQ71 | Chromosome 3 | ATP-binding | 15.60 |
| A0A1C3YLN5 | Chromosome 4 | - | 14.31 |
| A0A1C3YLW0 | Chromosome 4 | Membrane | 14.28 |
| I1RA25 | Chromosome 1 | Exocytosis | 14.50 |
| I1RC58 | 1-phosphatidylinositol 4-kinase (EC 2.7.1.67) | Kinase | 13.61 |
| I1RHY5 | Chromosome 2 | Disulfide bond | 13.60 |
| I1RI16 | Chromosome 2 | Heme | 15.09 |
| I1RIK7 | Chromosome 2 | - | 14.13 |
| I1RKJ4 | Chromosome 2 | - | 15.43 |
| I1RUN2 | Chromosome 4 | - | 15.31 |
| I1RXX8 | Chromosome 4 | Apoptosis | 15.59 |
| I1RYX5 | V-type proton ATPase subunit F | Hydrogen ion transport | 14.85 |
| I1S2H7 | Chromosome 3 | - | 15.00 |
| I1S2L2 | Chromosome 3 | - | 14.46 |
| Q4IBS9 | Vacuolar protein-sorting protein BRO1 | Coiled coil | 12.98 |
| V6RIE5 | Very-long-chain 3-oxoacyl-CoA reductase (EC 1.1.1.330) | Endoplasmic reticulum | 14.78 |
| V6RXE7 | choline-phosphate cytidylyltransferase (EC 2.7.7.15) | Lipid biosynthesis | 14.87 |
| A0A1C3YJK5 | cysteine desulfurase (EC 2.8.1.7) | Iron | 13.97 |
| A0A1C3YKF5 | Chromosome 3 | Hydrolase | 16.00 |
| A0A1C3YLU7 | Chromosome 4 | - | 17.03 |
| I1RE84 | Chitin synthase 5 (EC 2.4.1.16) | Actin-binding | 15.63 |
| I1RGI5 | Chromosome 2 | - | 15.78 |
| I1RI07 | Chromosome 2 | NADP | 16.18 |
| I1RMQ7 | Chromosome 3 | - | 15.57 |
| I1RPE1 | Chromosome 3 | Coiled coil | 14.02 |
| I1RYS0 | Chromosome 4 | ATP-binding | 15.02 |
| I1S4Y2 | Chromosome 1 | - | 15.47 |
| A0A098DCL3 | Chromosome 2 | - | 15.83 |
| A0A1C3YKY9 | hydroxyacylglutathione hydrolase (EC 3.1.2.6) (Glyoxalase II) | Hydrolase | 15.62 |
| I1R9D1 | Chromosome 1 | - | 15.07 |
| I1R9L9 | beta-glucosidase (EC 3.2.1.21) | Carbohydrate metabolism | 15.30 |
| I1RC80 | Chromosome 1 | - | 15.63 |
| I1RCY4 | Chromosome 1 | NADP | 14.76 |
| I1RGF9 | Chromosome 2 | - | 15.98 |
| I1RJP3 | Chromosome 2 | - | 16.92 |
| I1RNR7 | Chromosome 3 | Membrane | 15.51 |
| I1RRU6 | Chromosome 4 | Hydrolase | 15.96 |
| I1RUP2 | Protein SDS23 (Protein sds23) | CBS domain | 15.03 |
| I1RXH2 | Chromosome 4 | - | 15.41 |
| I1S104 | Longiborneol synthase CLM1 (EC 4.2.3.-) | 3D-structure | 15.56 |
| V6RGQ1 | Chromosome 4 | Membrane | 15.78 |
| A0A0E0SGL9 | Chromosome 4 | Membrane | 15.22 |
| A0A0E0SNT6 | Chromosome 3 | - | 13.93 |
| A0A1C3YLV7 | Chromosome 4 | Acyltransferase | 15.33 |
| I1RCS1 | Methionine aminopeptidase (EC 3.4.11.18) | Aminopeptidase | 14.70 |
| I1RDS5 | Chromosome 1 | Acyltransferase | 16.01 |
| I1REV6 | Chromosome 1 | - | 15.17 |
| I1RI19 | Chromosome 2 | Flavoprotein | 14.88 |
| I1RMQ3 | Chromosome 3 | - | 15.90 |
| I1RWA0 | Chromosome 2 | - | 14.08 |
| I1S0B0 | Chromosome 1 | - | 14.14 |
| I1S331 | Chromosome 3 | - | 17.14 |
| Q00909 | Trichodiene synthase (EC 4.2.3.6) | Lyase | 14.01 |
| A0A098DGK5 | Chromosome 2 | - | 15.58 |
| A0A098DGV9 | Chromosome 2 | - | 15.51 |
| A0A098DNT5 | Chromosome 2 | ANK repeat | 15.78 |
| A0A0E0SBD5 | Chromosome 4 | Membrane | 13.94 |
| I1RP53 | Cyanide hydratase (CHT) (EC 4.2.1.66) | Hydrolase | 15.57 |
| I1RTS7 | Beta-glucuronidase (EC 3.2.1.31) | Glycosidase | 15.61 |
| I1RUV7 | Chromosome 2 | Membrane | 15.65 |
| I1S4Q6 | Chromosome 1 | Hydrolase | 14.37 |
| V6R882 | Chromosome 2 | Membrane | 14.19 |
| A0A1C3YLZ2 | Chromosome 4 | NADP | 15.79 |
| I1RDU2 | Chromosome 1 | - | 15.40 |
| I1RF16 | Amine oxidase (EC 1.4.3.-) | Copper | 15.22 |
| I1RL06 | ZEB2-regulated ABC transporter 1 | ATP-binding | 14.35 |
| I1RLG1 | Chromosome 3 | Aspartyl protease | 15.39 |
| I1S2Y3 | Chromosome 3 | Oxidoreductase | 15.76 |
| A0A098DCT0 | Chromosome 2 | Heme | 15.54 |
| A0A1C3YIB0 | Chromosome 1 | - | 15.51 |
| I1R9D3 | Alpha-galactosidase (EC 3.2.1.22) | Glycosidase | 15.85 |
| I1R9P5 | Chromosome 1 | - | 15.16 |
| I1RGM2 | Chromosome 2 | - | 15.68 |
| I1RIS3 | Chromosome 2 | - | 15.50 |
| I1RT94 | Serine/threonine-protein phosphatase (EC 3.1.3.16) | Hydrolase | 14.36 |
| I1S1K3 | Chromosome 1 | FAD | 15.12 |
| I1S340 | Chromosome 3 | - | 15.63 |
| V6QT89 | nicotinate phosphoribosyltransferase (EC 6.3.4.21) | Ligase | 15.61 |
| A0A098DCN8 | Chromosome 2 | - | 16.10 |
| A0A098DTY1 | Chromosome 4 | Membrane | 14.47 |
| V6RUG6 | Chromosome 4 | Maltose metabolism | 14.33 |
| I1S1G0 | chitinase (EC 3.2.1.14) | Carbohydrate metabolism | 15.91 |
| I1S5G4 | Chromosome 1 | FAD | 16.15 |
| A0A098DXB4 | Trehalase (EC 3.2.1.28) | Glycosidase | 15.78 |
| I1RQ23 | Signal recognition particle subunit SRP68 (SRP68) | Cytoplasm | 14.17 |
| Q4ICI6 | Ceramide-binding protein SVF1 (Survival factor 1) | Cytoplasm | 15.88 |
| A0A098DHI5 | Chromosome 2 | - | 15.49 |
| A0A098DVZ3 | Structural maintenance of chromosomes protein | ATP-binding | 14.29 |
| I1RQ95 | Chromosome 3 | - | 15.99 |
| A0A098D104 | Chromosome 1 | - | 16.19 |
| A0A1C3YJM7 | Amine oxidase (EC 1.4.3.-) | Copper | 15.52 |
| I1RDN8 | Chromosome 1 | - | 16.58 |
| I1RGQ2 | Chromosome 2 | - | 13.59 |
| I1RJ32 | Chromosome 2 | Membrane | 16.30 |
| I1RIT7 | Chromosome 2 | - | 15.16 |
| I1RYZ9 | Chromosome 4 | - | 15.57 |
| I1RG33 | Chromosome 1 | Flavoprotein | 15.16 |
| I1RFZ7 | Chromosome 1 | DNA-binding | 13.72 |
| I1RVD8 | Highly reducing polyketide synthase PKS6 (HR-PKS PKS6) (EC 2.3.1.-) | Acyltransferase | 15.28 |
| A0A0E0S5W7 | Chromosome 2 | - | 15.98 |
| I1RF48 | Chromosome 1 | Carbohydrate metabolism | 16.43 |
| I1RTP0 | Chromosome 4 | - | 15.15 |
| I1RUD4 | Chromosome 4 | - | 15.02 |
| I1R9C2 | ubiquitinyl hydrolase 1 (EC 3.4.19.12) | Hydrolase | 14.39 |

*Protein ID from UniProt

^#^Keyword and function derived from UniProt terms.

**Table S5: List of hub proteins from core wheat proteome**

| **Protein ID^*^** | **Protein name** |
| --- | --- |
| W5I1R7_WHEAT | 30S ribosomal protein S3, chloroplastic |
| W5FEZ3_WHEAT | 40S ribosomal protein S12 |
| A0A1D6S1Y9 | 40S ribosomal protein S15 |
| A0A3B6LV48 | 40S ribosomal protein S15 |
| W5DS33_WHEAT | 40S ribosomal protein S16 |
| W5FFT4_WHEAT | 40S ribosomal protein S16 |
| A0A3B6IZD6 | 50S ribosomal protein L11, chloroplastic |
| A0A3B6RK63 | 60S acidic ribosomal protein P0 |
| A0A3B6LTN4 | Ribosomal protein L13a |
| RPL3 | Ribosomal protein L3 (Ribosomal protein L3-B3) (Ribosomal protein L3B-1) (Ribosomal protein L3B-3) |
| Q5I7K7_WHEAT | Ribosomal protein L39 |
| RPL3-A3 | Ribosomal protein L3-A3 (Ribosomal protein L3A-3) |
| RPL3-B2 | Ribosomal protein L3-B2 (Ribosomal protein L3B-2) |
| A0A3B6HQ09 | Small ribosomal subunit protein uS10 domain-containing protein |
| A0A3B6K9W5 | Small ribosomal subunit protein uS10 domain-containing protein |
| A0A3B6HQ62 | Small ribosomal subunit protein uS2 |
| rps2 | Small ribosomal subunit protein uS2c (30S ribosomal protein S2, chloroplastic) |
| A0A3B5Y1V7 | Acyl carrier protein |
| A0A3B6HVH5 | Acyl carrier protein |
| A0A3B6JG13 | Aldehyde dehydrogenase |
| A0A3B6KIN5 | aldehyde dehydrogenase (NAD(+)) (EC 1.2.1.3) |
| A0A3B6RGM5 | Aldehyde dehydrogenase domain-containing protein |
| A0A3B6RJ64 | Aldehyde dehydrogenase domain-containing protein |
| A0A3B6TGW7 | Aldehyde dehydrogenase domain-containing protein |
| A0A3B6QKK6 | Aldehyde dehydrogenase domain-containing protein |
| A0A3B6NNP8 | Citrate synthase |
| A0A3B5XW54 | Dihydrolipoyl dehydrogenase (EC 1.8.1.4) |
| A0A3B5XZQ6 | DSK2a |
| A0A3B6SI60 | Elongation factor 1-gamma 2 |
| A0A0C4BJJ8 | fumarate hydratase (EC 4.2.1.2) |
| W5F826_WHEAT | Proteasome subunit beta |
| A0A3B6NLV6 | Succinate-semialdehyde dehydrogenase (EC 1.2.1.24) |
| A0A3B6JIP4 | Thioredoxin domain-containing protein |
| A0A0C4BJ55 | thioredoxin-dependent peroxiredoxin (EC 1.11.1.24) |
| A0A3B5Z4F9 | t-SNARE coiled-coil homology domain-containing protein |
| A0A3B6KJN0 | Ubiquitin-like domain-containing protein |
| A0A3B6SEI0 | Uncharacterized protein |
| A0A3B5XUE5 | Uncharacterized protein |

*Protein ID from UniProt
